# STAT1 sets microglial neutral-lipid content independently of lipid-handling transcriptional programs

**DOI:** 10.64898/2026.08.15.745002

**Authors:** Adil El Mesaoudi, Josephine Marie Boldt Lundby, Noor De Jong, Yonglun Luo, Lin Lin, Dong Won Kim

## Abstract

Inflammatory activation and lipid remodeling are linked features of microglial states, but how inflammatory transcription factors shape microglial lipid handling is unclear. Here we show that STAT1 sets neutral-lipid content in microglia through a route not predicted by lipid-handling transcription. Acute STAT1 depletion in primary microglia lowered neutral-lipid content while lipid-uptake and lipid-storage programs were induced, and interferon-γ activation moved inflammatory transcription in the opposite direction yet lowered lipid content alike. Single-cell transcriptomic and chromatin profiling of Stat1- and Irf1-deficient mice showed that STAT1 and IRF1 organize overlapping inflammatory and lipid-handling programs, with genome-wide accessibility changes that did not predict transcriptional output at individual lipid-handling loci. Microglia co-expressing STAT1 and APOE recurred across Alzheimer’s disease and multiple sclerosis datasets. Transcriptional program engagement is therefore separable from cellular lipid state, and lipid-handling gene expression cannot be read as a proxy for microglial lipid content.

## Introduction

Microglia are the resident immune cells of the central nervous system, maintaining tissue homeostasis and coordinating responses to injury, infection, and neurodegeneration^1,2^. Under inflammatory or degenerative stress they transition into distinct cellular states through coordinated gene reprogramming^3,4^, and these transitions are accompanied by metabolic remodeling, including changes in lipid uptake, storage, and utilization^5–7^. Lipid droplets, which are intracellular organelles that store neutral lipids such as triglycerides and cholesteryl esters, accumulate in macrophages and microglia during infection, aging, and neurodegenerative disease, and their accumulation coincides with altered lipid metabolism, phagocytosis, and inflammatory output ^8–11^. Lipid handling is therefore a regulated component of activated microglial states rather than a passive consequence of lipid excess^12,13^.

Single-cell transcriptomic studies have identified multiple microglial states enriched for lipid-handling programs^1,14^, which commonly express *Apoe*, *Lpl*, and other genes involved in lipid uptake, trafficking, processing, and turnover^15,16^. Inflammatory signaling and lipid handling are directly coupled in at least some of these states. In the median eminence, an *Apoe*-positive microglial population engages lipid-handling, interferon, and stress-response programs, Western diet increases both interferon signaling and lipid accumulation in these cells, and microglial *Apoe* deletion attenuates the coupling^17^. The transcriptional regulators that coordinate inflammatory and lipid-handling programs in adult microglia nonetheless remain largely undefined.

STAT1 is a central regulator of cytokine and innate immune signaling and a best known transcription factor for activating interferon-stimulated genes that drive antimicrobial and inflammatory responses^18–21^. Its activity extends beyond canonical interferon-responsive transcription: STAT1 can reshape broader cellular programs and influence chromatin accessibility, positioning it to establish the regulatory context in which inflammatory responses occur^22,23^ Because inflammatory signaling and lipid handling are tightly coordinated in myeloid cells, STAT1 is well placed to connect inflammatory state with lipid-handling transcription^3,24^. Interferon regulatory factor 1 (IRF1) is induced downstream of STAT1 and mediates a substantial component of interferon-responsive transcription^18,25,26^, making the canonical STAT1-IRF1 hierarchy a natural framework for asking whether its output extends to lipid-handling programs. Other IRF-family members already show that interferon-associated factors can act on microglial identity and lipid biology: IRF8 configures postnatal microglial enhancer landscapes and transcriptional programs^27^, and IRF7 regulates microglial lipid-droplet accumulation after ischemic stroke through impaired lipophagy^28^. Whether STAT1 and IRF1 shape lipid-handling programs in adult microglia under basal in vivo conditions has not been systematically examined.

Here we show that STAT1 and IRF1 converge on inflammatory and lipid-handling programs in adult microglia, and that engaging these programs does not specify neutral-lipid content in a fixed direction. Combining single-cell transcriptomic and chromatin-accessibility profiling of *Stat1*- and *Irf1*-deficient mice with reanalysis of human and mouse disease datasets and acute STAT1 depletion in primary mouse microglia and human microglial-like cells, we find that chromatin accessibility, lipid-handling transcription, and neutral-lipid content can move independently of one another.

## Results

### STAT1 loss reshapes adult microglial states and reorganizes lipid-handling programs

To determine whether STAT1 shapes microglial transcriptional states under basal conditions, we performed single-cell RNA sequencing of cerebral cortical cells from 8-week-old wild-type (WT) and *Stat1*-deficient (Stat1^-/-^) mice (Fig. 1A). The dataset resolved the major neural, glial, vascular, and immune populations of the adult brain, each identified by canonical lineage markers (Fig. S1A, B). *Stat1* deficiency altered transcription in astrocytes, endothelial cells, and neurons, with distinct responses in each cell type (Fig. S1C-K; Tables S1-3).

**Figure 1.**
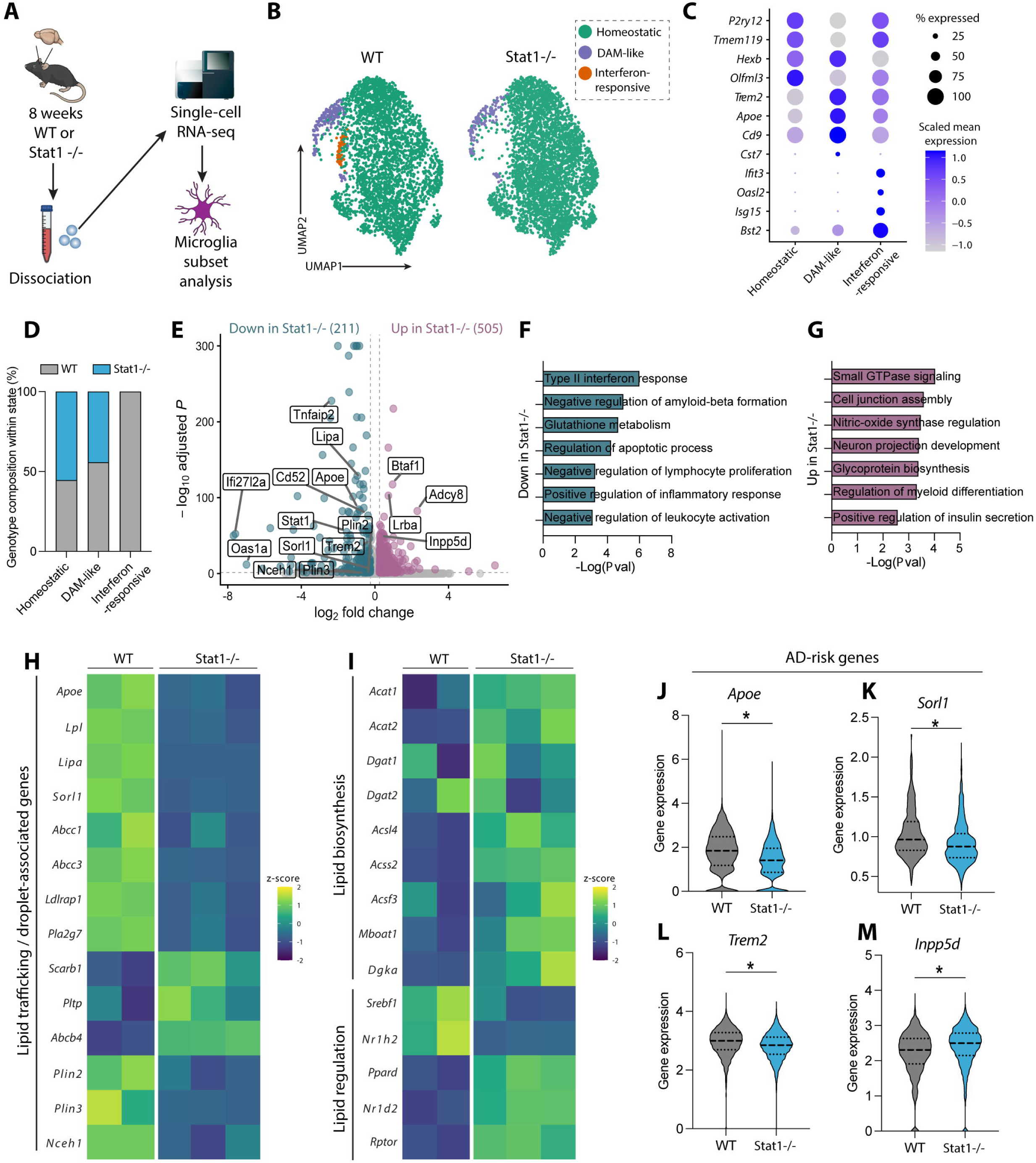
STAT1 loss reshapes adult microglial states and lipid-handling programs. **(A)** Experimental workflow. Cerebral cortices from 8-week-old wild-type (WT) and *Stat1*−/− mice were dissociated and analyzed by single-cell RNA sequencing (scRNA-seq), followed by isolation and analysis of the microglial compartment. **(B)** UMAP visualization of WT and *Stat1*−/− microglia colored by transcriptional state: homeostatic, disease-associated microglia (DAM)-like, and interferon-responsive. **(C)** Dot plot of representative marker genes used to annotate the three microglial states. Dot size indicates the percentage of cells expressing each gene, and color indicates scaled mean expression. **(D)** Proportions of homeostatic, DAM-like, and interferon-responsive microglia within each genotype. **(E)** Volcano plot of differential gene expression between *Stat1*−/− and WT microglia. Negative log fold-change values indicate lower expression in *Stat1*−/− microglia, positive values higher expression. Selected genes associated with inflammatory signaling, lipid handling, and microglial function are labeled. **(F, G)** Selected biological process terms enriched among genes expressed at lower (F) or higher (G) levels in *Stat1*−/− microglia relative to WT. Bar length indicates −log_10_(P value). **(H)** Heatmap of scaled expression of selected genes associated with lipid trafficking, lipid-droplet biology, cholesterol handling, and lysosomal lipid processing in WT and *Stat1*−/− microglia. Colors indicate row-scaled z-scores. **(I)** Heatmap of scaled expression of genes associated with lipid biosynthesis and lipid-regulatory programs in WT and *Stat1*−/− microglia. Colors indicate row-scaled z-scores. **(J-M)** Single-cell expression distributions of the Alzheimer’s disease risk-associated genes *Apoe* (J), *Sorl1* (K), *Trem2* (L), and *Inpp5d* (M) in WT and *Stat1*−/− microglia. Horizontal lines within the violins indicate the median and quartiles. *P < 0.05.

In microglia, unsupervised clustering resolved three states with homeostatic, disease-associated microglia (DAM)-like, and interferon-responsive features (Fig. 1B, C). Homeostatic microglia expressed *P2ry12*, *Tmem119*, *Hexb*, and *Olfml3*; the DAM-like population expressed *Trem2*, *Apoe*, *Cd9*, and *Cst7*; and a smaller interferon-responsive population expressed *Ifit3*, *Oasl2*, *Isg15*, and *Bst2* (Fig. 1C). *Stat1* deficiency reduced both the interferon-responsive and DAM-like fractions, with a corresponding increase in the homeostatic fraction (Fig. 1B, D). STAT1 therefore contributes to the organization of adult microglial states under basal conditions.

Differential-expression analysis of the microglial compartment identified 211 decreased and 505 increased genes in *Stat1*-deficient microglia (Fig. 1E; Table S4). Decreased genes were enriched for type II interferon responses, amyloid-beta formation, glutathione metabolism, and leukocyte activation (Fig. 1F). Increased genes were enriched for small GTPase signaling, cell-junction organization, glycoprotein biosynthesis, and myeloid differentiation (Fig. 1G). *Stat1* deficiency therefore reconfigures the adult microglial transcriptome beyond canonical interferon-responsive genes.

Many of these genes act in lipid handling. *Stat1*-deficient microglia showed reduced expression of *Apoe*, *Lpl*, *Lipa*, *Plin2*, *Plin3*, *Nceh1*, and *Sorl1*, which function in lipoprotein trafficking, lysosomal lipid processing, lipid-droplet organization, and cholesterol handling (Fig. 1H). Other lipid-handling genes, including *Scarb1*, *Pltp*, and *Abcb4*, increased (Fig. 1H). Genes involved in lipid biosynthesis and lipid-responsive regulation likewise changed in opposing directions (Fig. 1I).

These changes did not arise solely from the smaller interferon-responsive fraction. State-resolved analysis confirmed the expected decrease in *Ifit3*, *Oasl2*, *Isg15*, *Bst2*, and *Irf7*, but lipid-handling genes also changed within the homeostatic and DAM-like compartments, with magnitude and direction varying by gene and state (Fig. S2A). *Stat1* deficiency therefore alters transcription within established microglial states as well as their relative representation. *Stat1* deficiency also altered microglial genes at Alzheimer’s disease-associated loci: *Apoe*, *Sorl1*, and *Trem2* decreased, whereas *Inpp5d* increased (Fig. 1J-M, S2B).

Transcriptome-derived metabolic-task analysis predicted changes across lipid, amino-acid, carbohydrate, energy, and vitamin and cofactor metabolism (Fig. S2C-F). The largest lipid-associated effects were reduced phospholipid, sphingolipid, ceramide, and cardiolipin synthesis, alongside changes in glutathione and redox processes. These predictions reflect transcriptional potential and do not establish metabolite abundance or biochemical flux.

Together, these data show that *Stat1* deficiency reshapes adult microglial state organization and reorganizes lipid-handling transcription under basal in vivo conditions. Because individual lipid-handling genes change in opposite directions, STAT1 does not act on microglial lipid biology as a single uniform transcriptional module.

### STAT1 and IRF1 loss converge on inflammatory and lipid-handling programs in adult microglia

To determine whether the microglial phenotype of *Stat1*-deficient mice reflects a STAT1-specific function or a broader inflammatory transcription-factor network, we performed single-cell RNA sequencing of cerebral cortical cells from adult *Irf1*-deficient (*Irf1*-/-) mice to test whether STAT1 operates through its canonical downstream effector and integrated the microglial compartment with the WT and *Stat1*-deficient datasets (Fig. 2A). The *Irf1*-deficient dataset recovered the principal neural, glial, vascular, and immune populations, each identified by canonical lineage markers (Fig. S3A, B). *Irf1* deficiency also altered transcription in astrocytes, endothelial cells, and neurons (Fig. S3C-K; Tables S5-8).

**Figure 2.**
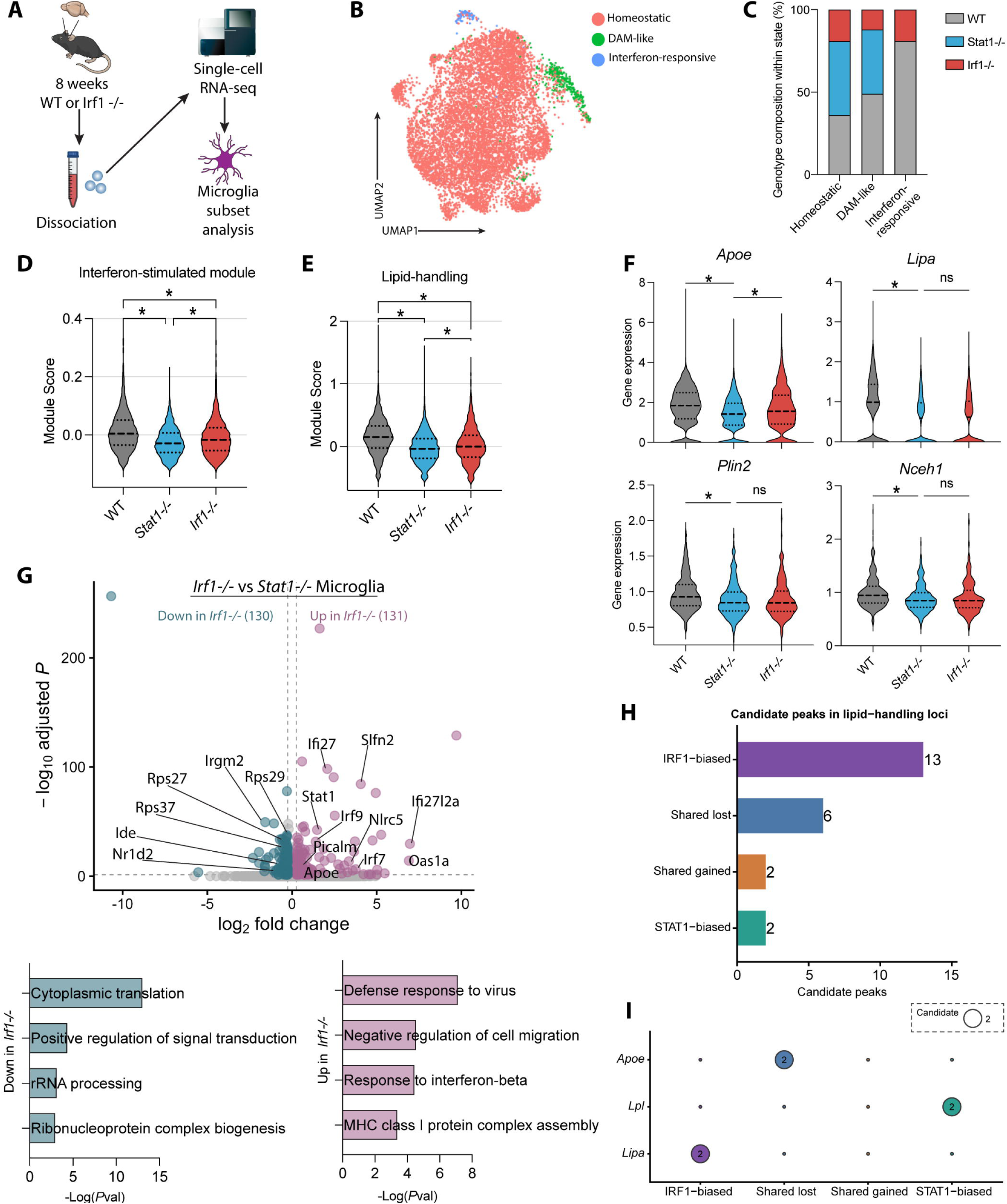
Loss of STAT1 or IRF1 produces overlapping inflammatory and lipid-handling responses and concordant chromatin-accessibility changes. **(A)** Experimental workflow. Cerebral cortices from 8-week-old WT and *Irf1*−/− mice were dissociated and analyzed by scRNA-seq. The microglial compartment was subsequently compared across WT, *Stat1*−/−, and *Irf1*−/− conditions. **(B)** Integrated UMAP visualization of WT, *Stat1*−/−, and *Irf1*−/− microglia colored by transcriptional state: homeostatic, DAM-like, and interferon-responsive. **(C)** Proportions of homeostatic, DAM-like, and interferon-responsive microglia within each genotype. The interferon-responsive state is absent in *Stat1*−/− and reduced but retained in *Irf1*−/− microglia. **(D, E)** Single-cell interferon-stimulated gene-module scores (D) and lipid-handling module scores (E) in WT, *Stat1*−/−, and *Irf1*−/− microglia. Horizontal lines within the violins indicate the median and quartiles. *P < 0.05. **(F)** Single-cell expression distributions of *Apoe*, *Lipa*, *Plin2*, and *Nceh1* in WT, *Stat1*−/−, and *Irf1*−/− microglia. *P < 0.05; ns, not significant. **(G)** Direct differential-expression analysis of *Irf1*−/− versus *Stat1*−/− microglia. Top, volcano plot of genes differentially expressed between the two knockout conditions; negative log_2_ fold-change values indicate lower expression in *Irf1*−/− relative to *Stat1*−/− microglia, positive values higher expression. Selected genes are labeled. Bottom, selected biological process terms enriched among genes expressed at higher or lower levels in *Irf1*−/− relative to *Stat1*−/− microglia. Bar length indicates −log_10_(P value). **(H)** Numbers of candidate differential-accessibility peaks within the analyzed lipid-handling loci, classified as IRF1-biased, shared lost, shared gained, or STAT1-biased according to their responses to *Irf1* or *Stat1* loss. Values at the ends of the bars indicate candidate-peak counts. **(I)** Distribution of candidate differential-accessibility peaks across the indicated lipid-handling loci and response classes. Bubble size indicates the number of candidate peaks assigned to each locus and response-class combination; values within larger bubbles indicate candidate-peak counts.

Integrated analysis resolved homeostatic, DAM-like, and interferon-responsive states across WT, *Stat1*-deficient, and *Irf1*-deficient microglia (Fig. 2B, C). Loss of either factor reduced the interferon-responsive and DAM-like fractions, with a corresponding increase in the homeostatic fraction (Fig. 2C). The two differed in degree: the interferon-responsive state was absent in *Stat1*-deficient microglia but reduced and retained in *Irf1*-deficient microglia (Fig. 2C). As expected for loss of two interferon-pathway factors, interferon-stimulated gene-module scores fell in both relative to WT (Fig. 2D). Lipid-handling module scores were also reduced in both (Fig. 2E).

At the gene level, *Stat1*-deficient and *Irf1*-deficient microglia showed reduced expression of *Lipa*, *Plin2*, and *Nceh1* relative to WT (Fig. 2F), spanning lysosomal lipid processing, lipid-droplet organization, and neutral-lipid turnover. Convergence was not uniform: *Apoe* decreased after loss of either factor, but more strongly in *Stat1*-deficient microglia (Fig. 2F).

Direct comparison of *Irf1*-deficient and *Stat1*-deficient microglia identified 130 genes expressed at lower and 131 at higher levels in *Irf1*-deficient cells (Fig. 2G; Table S9). Genes higher in *Irf1*-deficient microglia were enriched for antiviral defense, interferon-beta responses, and MHC class I complex assembly, indicating that STAT1 loss depletes interferon-responsive transcription more completely than IRF1 loss and matching the residual interferon-responsive state in *Irf1*-deficient microglia. Genes lower in *Irf1*-deficient microglia were enriched for cytoplasmic translation, signal transduction, rRNA processing, and ribonucleoprotein-complex biogenesis. STAT1 and IRF1 therefore produce strongly overlapping but nonidentical transcriptional outputs in adult microglia.

We next asked whether this convergence extended to chromatin accessibility. Integrated single-cell ATAC-seq showed broad overlap among the WT, *Stat1*-deficient, and *Irf1*-deficient microglial chromatin landscapes (Fig. S4A), and genome-wide accessibility effects were strongly correlated between the two (Fig. S4C). We classified candidate peaks as shared lost, shared gained, STAT1-biased, or IRF1-biased according to their responses to loss of each factor (Fig. S4B, D; Table S10).

Lipid-handling loci contained both shared and factor-biased peaks (Fig. 2H). *Apoe*, *Lpl*, and *Lipa* each carried a different combination of classes, so individual lipid-handling loci did not follow a uniform accessibility pattern (Fig. 2I).

Promoter and distal accessibility summaries reinforced this locus-level heterogeneity (Fig. S4E). Several genes with reduced RNA expression, including lipid-handling genes, showed increased promoter or distal accessibility in *Stat1*-deficient or *Irf1*-deficient microglia, so accessibility did not predict the direction of transcriptional change. Distal accessibility at lipid-handling loci trended higher than at matched control regions, but the difference did not survive correction for multiple testing and does not support selective opening of these loci as a class (Fig. S4F; Table S10).

Together, STAT1 and IRF1 converge on inflammatory and lipid-handling programs in adult microglia while retaining factor-biased outputs at selected genes. The mismatch between accessibility and expression argues against a model in which either factor controls lipid-handling genes through uniform opening or closing of individual loci.

### STAT1 APOE microglia recur across AD and MS disease contexts

To determine whether inflammatory and lipid-handling transcription intersect in disease-associated microglial states, we reanalyzed published mouse and human scRNA-seq datasets of Alzheimer’s disease (AD)^29,30^ and multiple sclerosis or experimental autoimmune encephalomyelitis (MS/EAE)^31^, asking whether microglia with detectable STAT1 and APOE transcripts recur across disease contexts and transcriptional states.

In the mouse AD dataset, microglia resolved into homeostatic, DAM1, DAM2, interferon-responsive, antigen-presenting, and disease-associated intermediate states (Fig. 3A, S5A). *Stat1* expression was concentrated in interferon-responsive microglia, whereas *Apoe*, *Lpl*, and *Plin2* were distributed across several activated and disease-associated states (Fig. 3B). Interferon-responsive microglia carried higher lipid-handling module scores than homeostatic microglia, indicating that inflammatory and lipid-handling transcription coexist within part of the disease-associated landscape (Fig. 3C).

**Figure 3.**
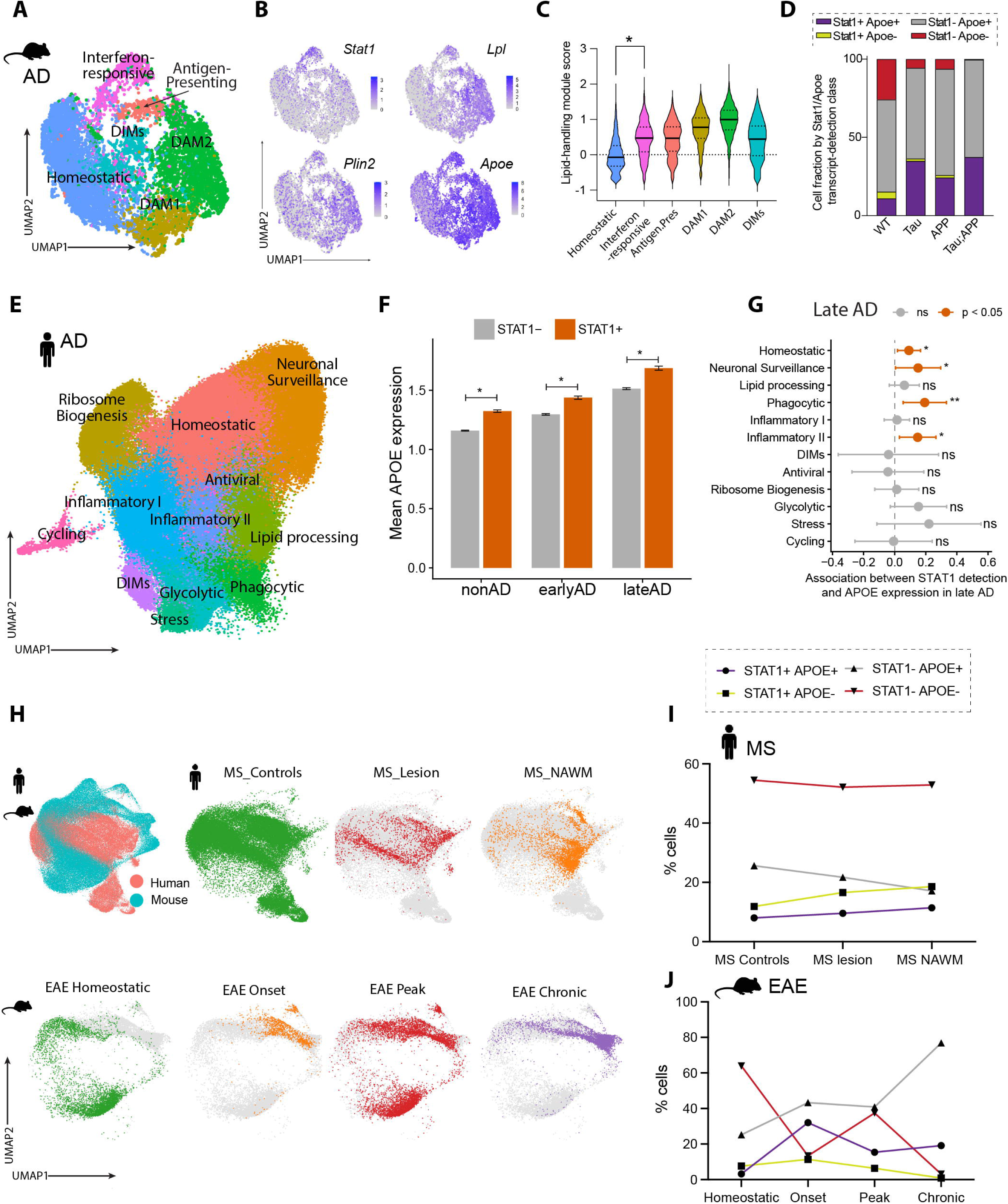
STAT1 APOE microglia recur across Alzheimer’s disease and multiple sclerosis contexts. **(A)** UMAP visualization of microglia from a published mouse Alzheimer’s disease (AD) scRNA-seq dataset, annotated as homeostatic, disease-associated microglia (DAM)1, DAM2, interferon-responsive, antigen-presenting, or disease-associated intermediate microglia (DIM). **(B)** Feature plots showing expression of *Stat1* and the lipid-handling genes *Lpl*, *Plin2*, and *Apoe* across the mouse AD microglial landscape. **(C)** Lipid-handling module scores across the indicated mouse microglial states. Horizontal lines within the violins indicate the median and quartiles; the dashed horizontal line indicates a module score of zero. *P < 0.05. **(D)** Relative proportions of microglia classified according to detectable *Stat1* and *Apoe* transcripts in WT, Tau, App, and combined Tau;App mouse models. Cells were classified as *Stat1*□*Apoe*□, *Stat1*□*Apoe*□, *Stat1*□*Apoe*□, or *Stat1*□*Apoe*□ according to transcript detection. **(E)** UMAP visualization of microglia from a published human AD single-nucleus RNA-seq dataset, annotated as homeostatic, neuronal-surveillance, ribosome-biogenesis, lipid-processing, phagocytic, Inflammatory I, Inflammatory II, disease-associated intermediate, antiviral, glycolytic, stress-associated, or cycling microglia. **(F)** Mean APOE expression in microglia with undetected or detected STAT1 transcript from non-AD, early AD, and late AD groups. Exact P values are indicated above the corresponding comparisons. **(G)** State-resolved association between STAT1 detection and APOE expression in late AD microglia. Points indicate estimated associations with STAT1 detection and horizontal lines indicate 95% confidence intervals. Positive estimates indicate higher APOE expression in cells with detectable STAT1 transcript. Significant associations are highlighted. *P < 0.05; **P < 0.01; ns, not significant. **(H)** UMAP visualization of mouse and human microglia from multiple sclerosis (MS) and experimental autoimmune encephalomyelitis (EAE) datasets. **(I)** Relative proportions of human microglia classified according to STAT1 and APOE transcript detection in MS control tissue, MS lesions, and normal-appearing white matter (NAWM). **(J)** Relative proportions of mouse EAE microglia classified as *Stat1 Apoe*, *Stat1 Apoe*, *Stat1 Apoe*, or *Stat1 Apoe* in naive animals and at disease onset, peak, and chronic stages.

Classifying cells by detectable *Stat1* and *Apoe* transcripts, *Stat1 Apoe* microglia were present in WT mice and rose in proportion in the Tau, APP, and combined Tau/APP models (Fig. 3D). The increase came primarily from a lower fraction of *Stat1 Apoe* cells, whereas *Stat1 Apoe* cells remained abundant and *Stat1 Apoe* cells rare across models. *Stat1* and *Apoe* co-detection therefore recurs across several mouse AD-associated genetic contexts rather than being restricted to one model.

In an independent human AD dataset spanning non-AD, early-AD, and late-AD samples (Fig. 3E, S5B), donor-adjusted analyses showed higher mean APOE expression in microglia with detectable STAT1 transcript than in STAT1-undetected cells, with the magnitude of the association varying among groups (Fig. 3F). Because this aggregate comparison can reflect differences in microglial state composition, we next examined the relationship within transcriptionally defined states.

Across all disease stages, detectable *STAT1* transcript was positively associated with *APOE* expression in several microglial states, with the strength of the association varying across the landscape (Fig. S5C). In Inflammatory II microglia, *STAT1* detection was associated with *APOE* but not with the other genes examined (Fig. S5D). In lipid-processing microglia, *STAT1* detection was associated with *APOE* and *LIPA* but not with *PPARG*, *ABCA1*, *TREM2*, or *LPL* (Fig. S5E). *STAT1* detection therefore tracked selected lipid-handling genes rather than a broad lipid-handling program.

Restricting the analysis to late AD gave a more state-selective pattern (Fig. 3G). *STAT1* detection was positively associated with *APOE* expression in homeostatic, neuronal-surveillance, phagocytic, and Inflammatory II microglia, but not in the other states examined. In late-AD Inflammatory II microglia, *STAT1* detection was also associated with higher *LIPA* expression (Fig. S5F), whereas in the late-AD lipid-processing population it was not associated with any tested lipid-handling gene (Fig. S5G). The relationship between *STAT1* and lipid-handling transcription therefore depends on both microglial state and disease stage.

We next asked whether *STAT1* and *APOE* co-expression also occurs during neuroinflammation. Integrated analysis of mouse EAE and human MS microglia identified shared and species-associated transcriptional structure (Fig. 3H). In human MS, *STAT1 APOE* microglia were present in control tissue, lesions, and normal-appearing white matter, with modest differences in pooled cell fractions among tissue categories (Fig. 3I). In mouse EAE, the pooled *Stat1 Apoe* fraction was highest at disease onset, fell at peak disease, and remained detectable in the chronic stage (Fig. 3J). Co-expression therefore extends beyond AD-associated states, and its abundance does not track disease severity monotonically.

Across a broader collection of published human and mouse microglial datasets, *STAT1* expression was highest in cytokine-responsive, interferon-associated, inflammatory, and MHC-associated superclusters in both species, and *STAT1 APOE* microglia were present in neurodegenerative, neuroinflammatory, vascular, and experimentally induced conditions (Fig. S6).

Together, *STAT1 APOE* microglia recur across independent mouse and human disease datasets. Their representation varies with genetic model and disease stage, and in human AD the association between detectable *STAT1* and lipid-handling genes, chiefly *APOE* and *LIPA*, is confined to particular microglial states. These analyses place the mouse genetic results in disease contexts in which inflammatory and lipid-handling transcription occupy the same cells.

### Acute STAT1 depletion lowers microglial neutral-lipid content while reorganizing lipid-handling transcription

Because constitutive knockouts reflect chronic loss, we asked whether STAT1 regulates microglial lipid content acutely and cell-intrinsically. We depleted *Stat1* in primary mouse microglia using siRNA (Fig. 4A). Of three *Stat1*-targeting siRNAs, siRNA #1 and siRNA #3 reduced *Stat1* expression most effectively and siRNA #2 less so (Fig. 4B); we carried siRNA #1 and #3 forward.

**Figure 4.**
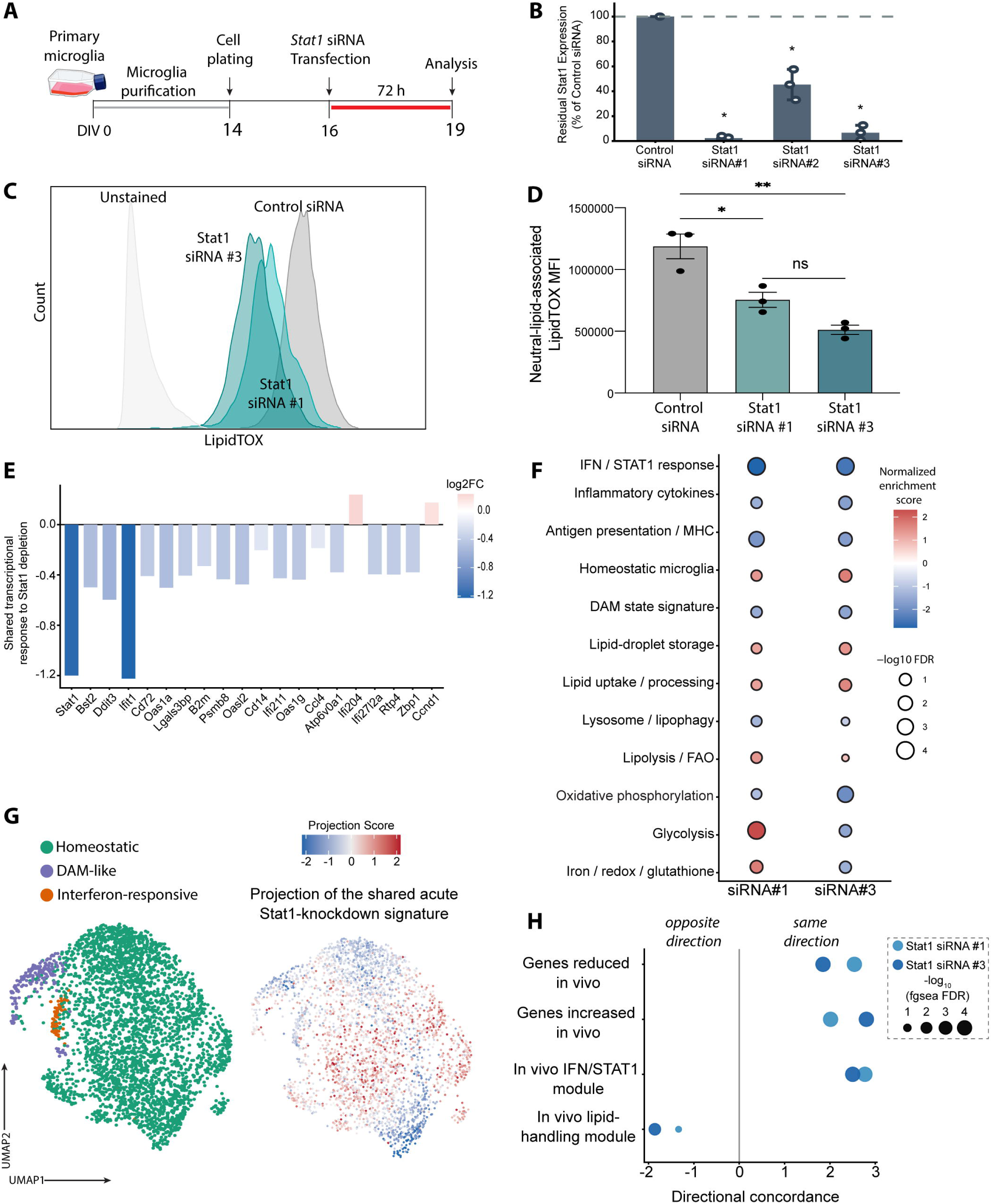
Acute STAT1 depletion lowers neutral-lipid content and partially reproduces the in vivo *Stat1*-loss response in primary microglia. **(A)** Experimental design for acute *Stat1* depletion in primary mouse microglia. Microglia were purified, plated on day in vitro (DIV) 14, transfected with *Stat1*-targeting siRNA at DIV16, and analyzed 72 h later at DIV19. **(B)** Residual *Stat1* expression following transfection with control siRNA or three independent *Stat1*-targeting siRNAs, shown relative to control-siRNA-transfected cells. Points represent independent replicates; bars indicate mean ± s.e.m. *P < 0.05. **(C)** Representative LipidTOX fluorescence distributions for control and *Stat1*-targeting siRNA conditions, with an unstained sample shown for reference. **(D)** Quantification of neutral-lipid content as LipidTOX median fluorescence intensity (MFI). Points represent independent replicates; bars indicate mean ± s.e.m. *P < 0.05; **P < 0.01; ns, not significant. **(E)** Selected genes showing concordant directional responses to *Stat1* siRNA #1 and #3. Bars indicate log fold change relative to control-siRNA-treated cells, and color indicates effect magnitude and direction. **(F)** Gene-set enrichment analysis of transcriptional responses to *Stat1* siRNA #1 and #3. Rows indicate predefined inflammatory, microglial-state, lipid-handling, metabolic, and redox-associated gene sets. Color indicates normalized enrichment score; bubble size indicates −log_10_ false-discovery rate (FDR). **(G)** Projection of the acute *Stat1*-knockdown transcriptional response onto the adult microglial landscape *in vivo.* Left, UMAP of adult microglia annotated as homeostatic, DAM-like, or interferon-responsive. Right, projection score indicating the relative similarity of individual in vivo microglia to the acute knockdown signature; positive and negative scores indicate similarity to opposing directions of the response. **(H)** Directional concordance between acute *Stat1* depletion in primary microglia and transcriptional programs defined in *Stat1*−/− adult microglia *in vivo*. Gene sets comprise genes expressed at lower or higher levels in *Stat1*−/− microglia, the in vivo IFN/STAT1 module, and the *in vivo* lipid-handling module. Positive values indicate concordance with the *in vivo* direction, negative values an inverse response. Bubble size indicates −log_10_ FDR; color distinguishes *Stat1* siRNA #1 and #3.

*Stat1* depletion did not affect baseline viability or microglial morphology (Fig. S7A, B), and LPS induced marked morphological activation in both control and depleted cultures, confirming that the cells remained responsive (Fig. S7A). Both *Stat1*-targeting siRNAs reduced neutral-lipid LipidTOX fluorescence relative to control siRNA (Fig. 4C, D; S7C). The same direction of change was obtained with BODIPY (data not shown). Acute STAT1 depletion therefore lowers neutral-lipid content in primary microglia without overt cytotoxicity.

Bulk RNA sequencing confirmed disruption of the STAT1-dependent program. Both siRNAs reduced *Stat1* and suppressed canonical interferon-responsive genes, including *Ifit1*, *Bst2*, *Oas1a*, *Oas1g*, *Oasl2*, *Lgals3bp*, *Psmb8*, *Rtp4*, and *Zbp1*, although a small number of genes, including *Ifi204* and *Ccnd1*, increased (Fig. 4E; S7D; Table S11). Broader transcriptional responses differed between the two reagents, so we restricted interpretation to responses shared by both, of which suppression of the STAT1 and interferon-response program was the core.

Gene-set enrichment analysis separated microglial state signatures from lipid-process gene sets (Fig. 4F). Both siRNAs suppressed the IFN/STAT1-response, inflammatory-cytokine, antigen-presentation, and DAM state signatures while increasing the homeostatic microglial signature. Lipid-process sets moved differently: lipid-droplet storage and lipid uptake or processing were positively enriched, lipolysis and fatty-acid oxidation trended positive, and lysosome or lipophagy sets were modestly negative. Oxidative phosphorylation decreased with both siRNAs, whereas glycolysis and iron, redox, and glutathione sets responded in opposite directions between the two reagents and were not interpreted further. Lipid-storage and lipid-uptake transcription therefore rose while neutral-lipid content fell.

Gene-level responses matched this split (Fig. S7D). Interferon-responsive genes decreased consistently with both siRNAs, whereas *Apoe*, *Lipa*, *Lpl*, *Plin2*, *Plin3*, *Pnpla2*, *Cpt1a*, *Abca1*, *Trem2*, and other lipid-handling genes responded heterogeneously. The neutral-lipid phenotype therefore did not follow uniform suppression of genes controlling lipid uptake, storage, processing, or utilization.

We next asked how closely acute depletion reproduced the transcriptional phenotype of *Stat1*-deficient adult microglia. Projection of the knockdown signatures onto the adult microglial landscape distributed the response across several regions of the state space rather than confining it to one state (Fig. 4G; S7E; S8A-F; Table S12). Gene-level effects correlated positively between acute depletion and constitutive *Stat1* loss, more strongly for siRNA #3 than for siRNA #1 (Fig. S8G, H). STAT1 and interferon-responsive genes formed the most conserved component of this cross-model response, whereas lipid-handling genes diverged between the in vivo and cultured systems.

Signature-level analysis gave the same result (Fig. 4H). Genes reduced or increased in *Stat1*-deficient adult microglia shifted in the same direction after acute knockdown, and the *in vivo* IFN/STAT1 module showed the strongest directional concordance. The *in vivo* lipid-handling module, by contrast, moved in the opposite direction after acute depletion with both siRNAs.

Together, acute STAT1 depletion lowers neutral-lipid content in primary microglia. Acute and constitutive *Stat1* loss share a conserved STAT1 and interferon-response program, whereas lipid-handling transcription differs between cultured and adult microglia. In neither system does the lipid phenotype follow uniform suppression of a single lipid-handling pathway.

### Neutral-lipid content is decoupled from inflammatory and PPAR**γ**-responsive transcription in STAT1-depleted microglia

Having established that acute STAT1 depletion reduces neutral-lipid content in primary microglia, we asked whether these cells retained transcriptional responsiveness to inflammatory and lipid-regulatory stimulation, using IFNγ and the PPARγ agonist pioglitazone respectively (Fig. 5A).

**Figure 5.**
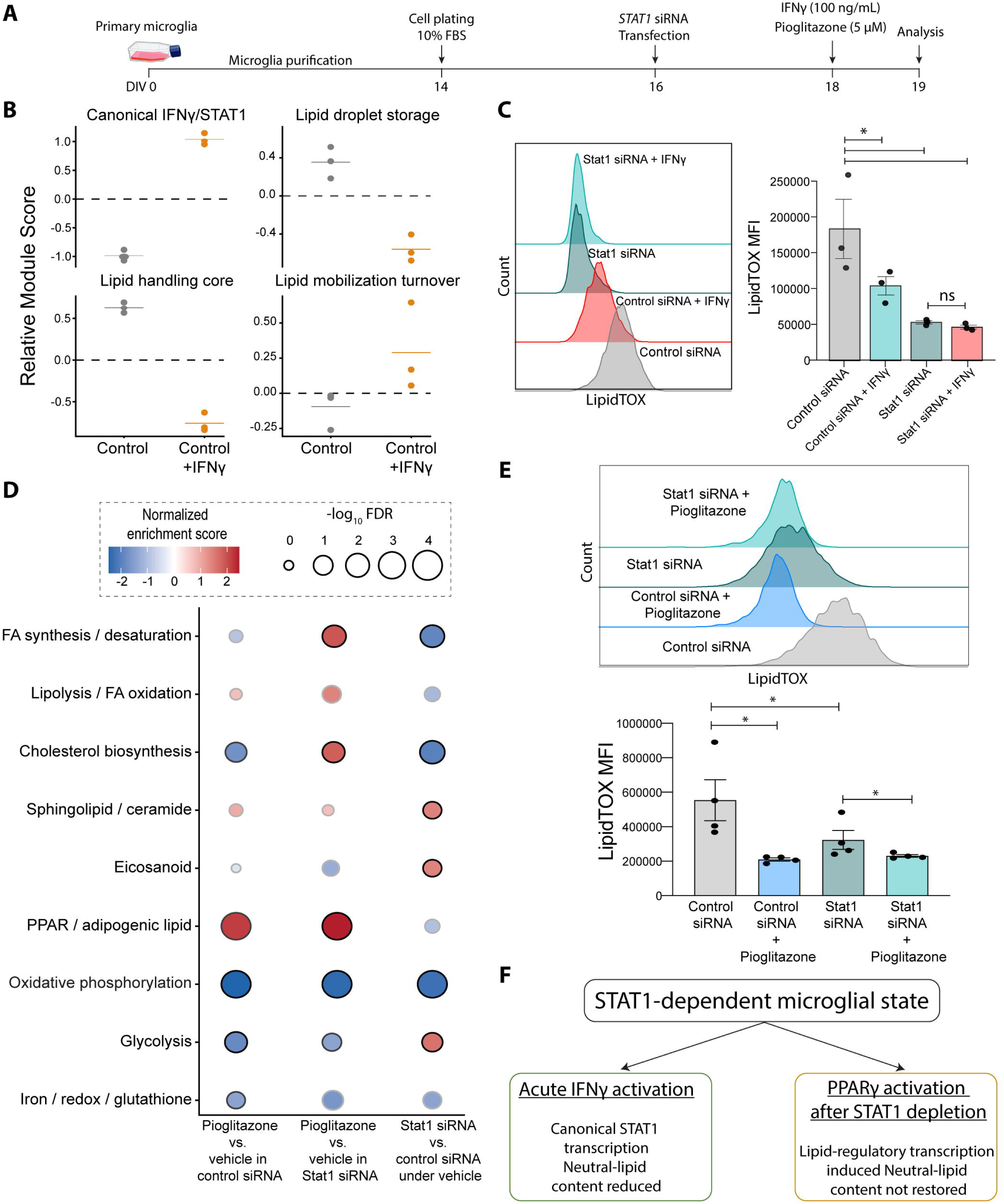
Inflammatory and lipid-regulatory transcription are uncoupled from neutral-lipid abundance in microglia. **(A)** Experimental design for IFNγ and pioglitazone perturbation in primary microglia. Microglia were purified from mixed glial cultures and plated at DIV14 in medium containing 10% FBS, transfected with control or *Stat1*-targeting siRNA at DIV16, treated at DIV18 with IFNγ or the PPARγ agonist pioglitazone, and analyzed at DIV19. **(B)** Transcriptional response of primary microglia to acute IFNγ stimulation. Relative module scores for canonical IFN/STAT1, lipid-droplet storage, core lipid handling, and lipid mobilization and turnover programs in control and IFNγ-treated cells. Points represent individual biological replicates. **(C)** Neutral-lipid content measured by LipidTOX staining and flow cytometry. Left, representative LipidTOX fluorescence distributions; right, quantification of LipidTOX median fluorescence intensity (MFI). Points represent independent biological replicates; bars indicate mean ± s.e.m. *P < 0.05; ns, not significant. **(D)** Gene-set enrichment analysis of transcriptional responses to pioglitazone and *Stat1* depletion. Columns show the pioglitazone response in control cells, the pioglitazone response in *Stat1*-depleted cells, and *Stat1* depletion alone relative to control. Rows indicate predefined metabolic, lipid-handling, and redox-associated programs. Color indicates normalized enrichment score; bubble size indicates −log_10_ false-discovery rate (FDR). **(E)** LipidTOX MFI following pioglitazone treatment in control and *Stat1*-depleted primary microglia. Left, representative LipidTOX fluorescence distributions; right, quantification of LipidTOX MFI. Points represent independent biological replicates; bars indicate mean ± s.e.m. *P < 0.05. **(F)** Proposed model. STAT1-depleted primary microglia retain a pioglitazone-induced PPARγ-responsive transcriptional program despite persistently reduced neutral-lipid content, and IFNγ activates inflammatory transcription while also reducing neutral-lipid content. Together these perturbations indicate that inflammatory or lipid-regulatory transcriptional activation does not predict the direction of change in neutral-lipid abundance.

IFNγ strongly induced *Stat1*, *Irf1*, *Isg15*, *Ifit1*, *Ifit2*, *Ifit3*, *Rsad2*, *Cxcl10*, *Socs1*, and MHC class II-associated genes, confirming robust engagement of the canonical inflammatory response (Fig. 5B; S9A; Table S13). IFNγ simultaneously reduced genes involved in lipid trafficking, lipid-droplet organization, cholesterol handling, and lysosomal lipid processing, including *Apoe*, *Lpl*, *Lipa*, *Plin2*, *Plin3*, *Soat1*, *Abca1*, *Cd36*, *Lrp1*, *Pparg*, and *Trem2*. Module-level analysis showed the same split: induction of the IFNγ and STAT1 program alongside suppression of the core lipid-handling and lipid-droplet-storage programs (Fig. 5B, S9A).

IFNγ also reduced neutral-lipid LipidTOX fluorescence in control microglia (Fig. 5C, S9B). STAT1 depletion lowered LipidTOX fluorescence on its own, and IFNγ produced little further change in depleted cells; in both conditions the signal remained well above the unstained control (Fig. 4C). Inflammatory activation and STAT1 depletion therefore both reduced neutral-lipid content while moving canonical STAT1-responsive transcription in opposite directions.

We next asked whether STAT1-depleted microglia retained the capacity to engage a defined lipid-regulatory pathway. PPARγ controls genes for fatty-acid uptake, lipid-droplet organization, lipid remodeling, triglyceride metabolism, and fatty-acid utilization, and *Pparg* itself was suppressed by IFNγ. We therefore treated control and STAT1-depleted microglia with pioglitazone (Fig. 5A).

Pioglitazone engaged lipid-regulatory transcription in both conditions (Fig. 5D; Tables S14, S15). In control microglia the response was dominated by extracellular lipid acquisition, trafficking, and processing: *Cd36* was most strongly induced, with positive responses in *Lpl*, *Plin2*, *Abcg1*, *Abca1*, and *Slc27a1*, whereas de novo fatty-acid synthesis genes including *Fasn*, *Acaca*, *Scd1*, and *Elovl6* did not increase coordinately. STAT1-depleted microglia again induced *Cd36* and *Plin2*, together with *Scd1*, *Dgat1*, *Lpin1*, *Lpcat3*, *Pnpla2*, and *Cpt1a*, and the coordinated PPARγ and adipogenic-lipid program remained inducible. STAT1 depletion therefore did not eliminate PPARγ-driven lipid-handling transcription.

This transcriptional induction did not translate into lipid accumulation. Pioglitazone reduced LipidTOX fluorescence in control cells and did not raise it in STAT1-depleted cells (Fig. 5E). Pioglitazone therefore reproduced, through an independent perturbation, the dissociation seen after STAT1 depletion: induction of lipid-uptake and lipid-droplet transcription alongside reduced neutral-lipid content.

We next tested whether increased lipid oxidation accounted for the reduced neutral-lipid content. The fluorescent sensor C11-BODIPY showed no selective increase in lipid peroxidation in STAT1-depleted microglia relative to controls (Fig. S9C), and pharmacological inhibition of lipid peroxidation with ferrostatin-1 did not restore neutral-lipid fluorescence (Fig. S9D). Generalized lipid peroxidation therefore does not account for the reduced LipidTOX signal under the conditions tested.

Limited lipid supply does not account for it either. Under serum-replete conditions, oleate supplementation did not reverse the reduced LipidTOX signal after STAT1 depletion (Fig. S9E). The assay nonetheless retained upward dynamic range: after serum starvation, oleate raised LipidTOX fluorescence dose-dependently at 20 and 250 µg/ml (Fig. S9F), indicating that serum-replete cultures were already lipid-loaded rather than unable to accumulate neutral lipids.

Together, STAT1-depleted primary microglia remain transcriptionally responsive to inflammatory and lipid-regulatory stimulation while their neutral-lipid content stays low (Fig. 5F). IFNγ induced canonical inflammatory transcription and lowered neutral lipid, whereas pioglitazone induced a coordinated PPARγ-responsive program without raising it. Neutral-lipid abundance in these cells is therefore not a simple readout of inflammatory or PPARγ-driven transcription, and its reduction after STAT1 depletion is explained neither by lipid peroxidation nor by limited lipid supply.

### STAT1 depletion increases neutral-lipid content in HMC3 cells, revealing model-dependent lipid outcome

To determine whether the neutral-lipid phenotype of primary mouse microglia generalizes to a human cell model, we turned to HMC3 microglial-like cells. LPS stimulation induced a robust inflammatory and antiviral transcriptional response, establishing HMC3 cells as a stimulus-responsive human system (Fig. 6A, B; S10; Tables S16, S17).

**Figure 6.**
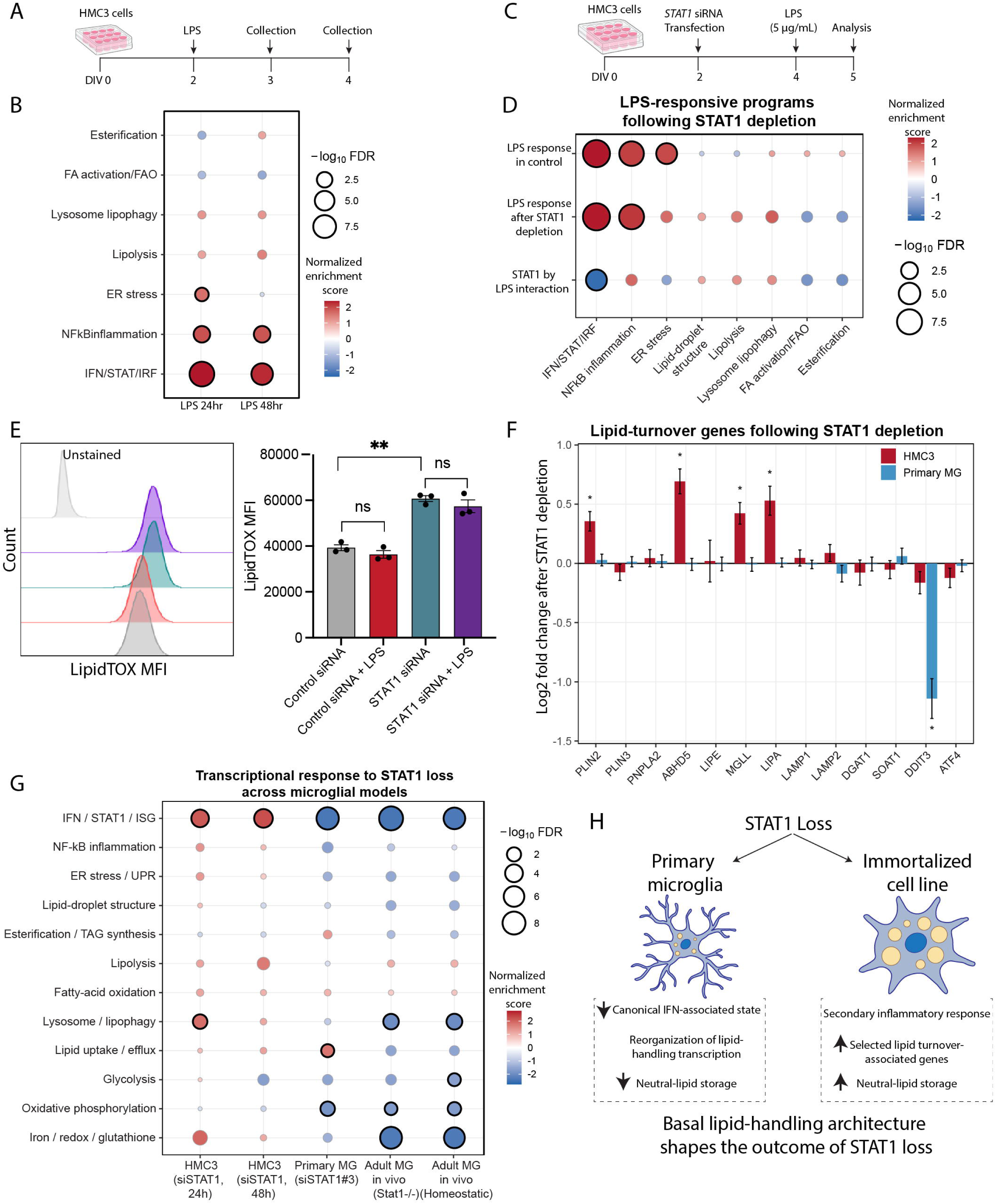
STAT1 depletion produces a model-dependent neutral-lipid phenotype in HMC3 cells. **(A)** Experimental design used to characterize the inflammatory response of HMC3 human microglial-like cells. Cells were treated with LPS at DIV2 and collected 24 or 48 h later for transcriptional analysis. **(B)** Gene-set enrichment analysis of transcriptional responses to LPS after 24 and 48 h. Rows indicate predefined inflammatory, stress, and lipid-handling programs, including IFN/STAT1/IRF signaling, NF-êB-associated inflammation, endoplasmic-reticulum stress, lipolysis, lysosome and lipophagy, fatty-acid activation and oxidation, and esterification. Color indicates normalized enrichment score (NES); bubble size indicates −log_10_ FDR. **(C)** Experimental design for analysis of the LPS response following acute STAT1 depletion. HMC3 cells were transfected with STAT1-targeting siRNA at DIV2, treated with LPS at DIV4, and analyzed at DIV5. **(D)** Gene-set enrichment analysis of the LPS response in control and STAT1-depleted HMC3 cells. Columns show the LPS response in control cells, the LPS response following STAT1 depletion, and the STAT1-by-LPS interaction. Rows indicate the predefined transcriptional programs. Color indicates NES; bubble size indicates −log_10_ FDR. **(E)** Neutral-lipid content measured by LipidTOX staining and flow cytometry in control- and STAT1-siRNA-treated HMC3 cells in the presence or absence of LPS. Left, representative LipidTOX fluorescence distributions, with an unstained sample shown for reference. Right, LipidTOX mean fluorescence intensity (MFI). Points represent independent replicates; bars indicate mean ± s.e.m. **P < 0.01; ns, not significant. **(F)** Gene-level changes in lipid-handling and stress-response genes following STAT1 depletion in HMC3 cells and primary microglia. Bars indicate shrunken log_2_ fold changes relative to the corresponding STAT1-intact control; positive values indicate higher expression following STAT1 depletion, negative values lower expression. Error bars indicate the standard error of the shrunken estimate. *P < 0.05. **(G)** Comparative gene-set enrichment analysis of the response to STAT1 loss across HMC3 cells, primary microglia, and adult microglia *in vivo*. Columns show HMC3 cells at 24 and 48 h after STAT1 depletion, primary microglia following acute *Stat1* depletion, all *Stat1*−/− microglia *in vivo*, and the homeostatic *Stat1*−/− subset. Rows indicate inflammatory, lipid-handling, metabolic, and redox-associated programs. Color indicates NES; bubble size indicates −log_10_ FDR. **(H)** Conceptual model summarizing the model-dependent effects of STAT1 loss on neutral-lipid homeostasis. In primary microglia, *Stat1* depletion suppresses canonical interferon-associated transcription, reorganizes lipid-handling programs, and reduces neutral-lipid content. In HMC3 cells, STAT1 depletion elicits a secondary inflammatory response, increases selected lipid-droplet and turnover-associated transcripts, and increases neutral-lipid content.

We then depleted STAT1 using two independent siRNAs, which reduced STAT1 expression relative to a non-targeting control without compromising cell viability (Fig. S11, S12), and treated control and STAT1-depleted cells with LPS (Fig. 6C). LPS-responsive transcription was broadly retained after depletion, with a restricted set of genes showing a significant STAT1-by-LPS interaction (Fig. 6D; S13A, B).

STAT1 depletion increased neutral-lipid LipidTOX fluorescence, both in the presence and absence of LPS (Fig. 6E; Tables S18, S19).

Transcriptional profiling after depletion showed that neither siRNA elicited a coordinated ER-stress response, whereas both elevated selected interferon- and lipid-turnover transcripts, including *IRF1*, *ISG15*, *IFIT1*, *MX1*, *PLIN2*, *ABHD5*, *MGLL*, and *LIPA* (Fig. 6F; S12; Tables S18, S19). STAT1 depletion in HMC3 cells therefore raised interferon-associated transcription, the opposite of its effect in primary mouse microglia.

Basal expression analysis showed that HMC3 cells expressed low or undetectable levels of several microglia-enriched lipid uptake and trafficking genes, including *APOE*, *TREM2*, *CD36*, and *MSR1*, while retaining core lipid-droplet, lipolytic, and lysosomal machinery (Fig. S13C).

STAT1 re-expression did not normalize the knockdown-associated LipidTOX phenotype, and STAT1 overexpression altered the signal in control cells (Fig. S14). These results preclude a clean rescue and indicate that the phenotype is sensitive to STAT1 abundance in both directions.

Comparative gene-set enrichment analysis across HMC3 cells, primary microglia, and adult microglia in vivo showed that the direction and magnitude of the response to STAT1 loss varied across systems, particularly for interferon-associated, lysosomal, oxidative-phosphorylation, glycolytic, and redox-associated programs (Fig. 6G). STAT1 depletion therefore has opposite effects on neutral-lipid fluorescence in HMC3 cells and primary mouse microglia (Fig. 6H). Because the two systems differ in baseline expression of microglia-enriched lipid uptake and trafficking genes and in their transcriptional response to STAT1 loss, this contrast marks a boundary on the transferability of the phenotype across cellular models rather than establishing a species difference.

## Discussion

We identify STAT1 and IRF1 as overlapping regulators of the transcriptional programs that organize adult microglial states and lipid handling, and we find that engagement of those programs does not specify neutral-lipid content in a fixed direction. Loss of either factor in vivo reshaped microglial states and reorganized lipid-handling transcription, with concordant genome-wide accessibility changes that nonetheless failed to predict transcriptional output at individual lipid-handling loci. In primary microglia, acute STAT1 depletion lowered neutral-lipid content while lipid-storage and lipid-uptake programs were positively enriched, and IFNγ activation lowered it in the same direction while moving inflammatory transcription the opposite way. HMC3 cells showed the reverse lipid phenotype. Together these results place STAT1 and IRF1 within a context-dependent framework linking inflammatory state, lipid-handling transcription, and neutral-lipid content.

Lipid droplets are dynamic organelles that integrate neutral-lipid storage with lipid mobilization, membrane homeostasis, and cellular stress responses^32,33^. In inflammatory myeloid cells, droplet formation and turnover are coupled to inflammatory signaling, and perturbing triglyceride storage or mobilization alters immune output^10,34,35^. Lipid-droplet-rich microglial states arise during aging and neurodegenerative disease, although their functional consequences vary across cellular and pathological contexts^8,9,11^. Neutral-lipid abundance is therefore a regulated cellular property rather than a passive consequence of lipid excess.

*In vivo*, *Stat1* deficiency altered the representation of homeostatic, DAM-like, and interferon-responsive states and moved lipid-handling genes in both directions, reducing a group involved in lipoprotein trafficking, lysosomal lipid processing, cholesterol handling, and lipid-droplet organization while increasing others. STAT1 therefore does not uniformly activate or suppress lipid-handling transcription; it reorganizes distinct components of the program. These changes were not attributable solely to the reduced interferon-responsive fraction, since lipid-handling genes also changed within the homeostatic and DAM-like compartments. Lipid-handling transcription is thus integrated with microglial state organization rather than controlled as a single uniform metabolic module.

Canonical models position STAT1 upstream of IRF1 in an interferon-responsive transcriptional cascade^36,37^. Our results are compatible with that hierarchy and extend its functional output to microglial lipid-handling transcription: loss of either factor reduced interferon-associated and lipid-handling module expression, and several lipid-handling genes responded similarly in the two genotypes. The outputs were not identical, however. *Apoe* expression fell after loss of either factor but more strongly after *Stat1* loss. Because state representation also differed between genotypes, this aggregate comparison reflects both compositional and within-state effects, and it does not establish direct regulation of the *Apoe* locus or a STAT1-exclusive mechanism. Individual genes may depend differently on STAT1, IRF1, other IRF-family factors, cooperating transcription factors, cofactor availability, and the underlying microglial state.

Chromatin accessibility adds a further regulatory dimension. Accessibility changes were concordant genome-wide between the two knockouts, yet individual lipid-handling loci carried different combinations of shared and factor-biased candidate peaks, and several genes with reduced transcript abundance showed increased promoter or distal accessibility. Accessibility alone therefore did not explain the observed transcriptional responses, consistent with contributions from transcription-factor occupancy, cofactor recruitment, enhancer-promoter communication, polymerase engagement, and three-dimensional chromatin organization. The candidate regions identified here provide a basis for subsequent direct-binding and locus-specific perturbation studies but do not themselves establish direct STAT1 or IRF1 regulation.

*Apoe* may have functional relevance beyond its use as a transcriptional marker. In median eminence microglia, microglial *Apoe* contributes to the relationship between interferon signaling and lipid accumulation during dietary stress, and *Apoe* deletion attenuates this coupling^17^. Changes in *Apoe* expression could therefore modify inflammatory and lipid-handling phenotypes rather than merely report a change in microglial state, although the present data do not test this directly. Any such consequence would also depend on APOE abundance, isoform, secretion, and lipidation state, none of which was measured here; a recent human microglial preprint reported distinct effects of lipid-bound and lipid-free APOE4 on lipid-droplet, mitochondrial, and interferon-associated phenotypes^38^.

The involvement of inflammatory transcription factors in microglial lipid regulation may extend beyond STAT1 and IRF1. IRF7 promotes lipid-droplet accumulation after ischemic stroke by impairing lipophagy through a *Gnai2*- and phosphatidylcholine-associated mechanism^28^, and IRF8 contributes to the configuration of microglial enhancer landscapes and transcriptional identity^27^. Individual interferon-associated transcription factors may therefore influence distinct aspects of microglial identity, lipid turnover, and inflammatory state. Whether STAT1 or IRF1 affects lipophagy, phospholipid remodeling, or related pathways in the present models remains unresolved.

The disease datasets provide an independent context in which inflammatory and lipid-handling transcription coexist. STAT1 and APOE co-detection recurred across mouse and human Alzheimer’s disease, EAE, and multiple sclerosis data, with context-dependent differences in representation across genetic models and disease stages. In human Alzheimer’s disease the association between detectable STAT1 and APOE varied by microglial state and disease stage rather than defining one invariant double-positive population. These analyses do not establish that STAT1 directly regulates APOE in human disease, that double-positive cells represent a distinct stable subtype, or that their abundance predicts disease progression. Such cells were also present in control contexts, and absolute proportions cannot be compared directly among datasets differing in tissue, disease stage, sequencing depth, and transcript-detection sensitivity. Their recurrence nonetheless supports the broader relevance of microglial states in which the two programs coexist.

Acute STAT1 depletion in primary microglia reproduced the canonical inflammatory component of the constitutive phenotype but not its lipid component. The lipid-handling module reduced in adult *Stat1*-deficient microglia was not coordinately suppressed in culture; instead, lipid-droplet-storage and lipid-uptake gene sets were positively enriched even as neutral-lipid-associated fluorescence fell. The functional lipid phenotype therefore could not be inferred from the direction of a single transcript, pathway, or gene module. One interpretation is that reduced neutral-lipid abundance elicits compensatory induction of lipid uptake and storage programs, although the present data do not establish this mechanism. Because LipidTOX reports neutral-lipid-associated fluorescence rather than lipid composition, flux, or intrinsic storage capacity, the discordance may also reflect a gap between pathway-level transcriptional potential and the specific neutral-lipid pool the assay detects.

The stimulation experiments sharpened this dissociation. IFNγ induced canonical STAT1-responsive, interferon-responsive, and antigen-presentation genes while suppressing multiple lipid-handling genes and reducing neutral-lipid-associated fluorescence. STAT1 depletion lowered neutral-lipid content as well, while suppressing rather than activating the same transcriptional program. Opposing effects on inflammatory transcription were therefore associated with the same direction of change in neutral-lipid abundance. This distinguishes acute cytokine stimulation from the basal STAT1-associated state of unstimulated microglia: STAT1 participates in inducible cytokine signaling, but its presence under basal conditions may also contribute to the broader transcriptional and cellular organization of microglia, so activation and loss need not produce reciprocal lipid outcomes. The present data do not identify the molecular features distinguishing these states, but they show that the magnitude or direction of canonical STAT1-responsive transcription alone does not predict neutral-lipid abundance.

PPARγ activation tested directly whether engagement of a defined lipid-regulatory pathway is accompanied by neutral-lipid accumulation. Pioglitazone induced genes involved in extracellular lipid acquisition, trafficking, lipid-droplet organization, phospholipid remodeling, triglyceride metabolism, lipid mobilization, and fatty-acid utilization, and STAT1-depleted microglia retained substantial responsiveness. Pioglitazone nonetheless did not increase neutral-lipid-associated fluorescence in either condition and reduced the signal in control cells. Because the agonist did not increase neutral-lipid content even in control microglia, this experiment cannot establish that STAT1 is required to translate PPARγ-responsive transcription into neutral-lipid storage. The supported conclusion is narrow: STAT1-depleted microglia remain transcriptionally responsive to PPARγ activation despite persistently reduced neutral-lipid-associated fluorescence.

Generalized lipid peroxidation does not appear to explain the reduced phenotype. STAT1 depletion did not selectively increase the C11-BODIPY oxidation signal, and ferrostatin-1 did not restore neutral-lipid-associated fluorescence. These findings do not exclude localized, transient, or lipid-species-specific oxidative changes, but they argue against a simple model in which generalized lipid peroxidation is the principal cause of the reduced LipidTOX signal.

The opposing neutral-lipid response in HMC3 cells defines the limits of transferability across cellular models. STAT1 depletion there increased neutral-lipid-associated fluorescence with two independent siRNAs and without reduced viability, and it increased rather than uniformly suppressed selected interferon-associated transcripts. The two systems differ in basal regulatory and lipid-handling architecture: HMC3 cells express very low levels of microglia-enriched lipid uptake and trafficking genes, including *APOE*, *LPL*, *TREM2*, *CD36*, and *MSR1*, while retaining broadly expressed lipid-droplet, lipolytic, and lysosomal machinery, and as an immortalized microglial-like line they do not reproduce the full transcriptional identity of primary human microglia^39,40^. The comparison should therefore not be interpreted as evidence of a mouse-versus-human species difference, but as evidence that the consequence of STAT1 depletion depends on the cellular system in which it occurs. The distinct baseline architectures are consistent with, but do not establish, a role for the underlying cellular state in determining the opposing lipid outcomes.

Several limitations define the next steps. The global *Stat1* and *Irf1* knockouts do not establish whether the in vivo effects are microglia-autonomous, since altered signaling in other brain or peripheral populations could influence microglial state, and because both models are germline nulls the adult phenotypes may also reflect developmental compensation and lifelong alterations in the tissue environment. Microglia-specific conditional perturbation will be required to resolve cell autonomy. The study does not establish direct STAT1 or IRF1 occupancy at lipid-handling regulatory elements; direct-binding assays and locus-specific perturbation will be necessary to distinguish direct targets from secondary responses to altered cellular state. The functional assays measure neutral-lipid-associated fluorescence but do not determine lipid composition, biochemical abundance, flux, or fate, or whether the detected pool consists predominantly of triglycerides, cholesteryl esters, or other species. Lipidomics, isotope tracing, and targeted manipulation of lipid uptake, esterification, lipolysis, lysosomal turnover, oxidation, and efflux will be required to identify the processes responsible for the reduced phenotype in primary microglia and the increased phenotype in HMC3 cells. Finally, differences between cultured and adult microglia indicate that culture conditions may modify the regulatory relationships identified here.

Together, our findings identify STAT1 and IRF1 as contributors to overlapping inflammatory and lipid-handling programs in adult microglia, extending the functional output of the canonical STAT1-IRF1 hierarchy to microglial lipid management. At individual lipid-handling loci, chromatin accessibility does not consistently predict transcriptional output, and across the cultured microglial models, engagement of inflammatory or lipid-regulatory transcription did not predict neutral-lipid abundance in a fixed direction. STAT1 perturbation therefore reveals a context-dependent separation between transcriptional state and neutral-lipid homeostasis.

## Methods

## 1. Mice

Homozygous *Stat1* knockout mice (B6.129S(Cg)-*Stat1tm1Dlv*/J, JAX #012606) and *Irf1* knockout mice (B6.129S2-*Irf1tm1Mak*/J, JAX #002762) were used for experiments. Age-matched wild-type (WT) mice served as controls. Female mice were used at 8 weeks of age. For tissue collection, whole cerebral cortices were rapidly dissected bilaterally, excluding non-cortical brain regions, and immediately processed for single-cell suspension preparation. Myelin-rich debris was reduced during subsequent debris-removal and density-gradient cleanup steps^29^.

Primary mouse microglia were prepared from postnatal day 1-2 C57BL/6JR pups obtained from time-mated females purchased from Janvier Labs.

## 2. Single-cell Sequencing

### 2.1 scRNA-seq cell preparation with transcriptional inhibition

Cells were isolated using a modified cortical dissociation protocol^29^ incorporating transcriptional and translational inhibitors to minimize activity-dependent artifacts in microglia during tissue processing, adapted from^41^.

Inhibitors were prepared fresh, and all solutions were maintained on ice and protected from light during tissue handling. Actinomycin D (Cat. No. A1410) was dissolved in DMSO at 5 mg/mL, triptolide (Cat. No. T3652) at 10 mM, and anisomycin (Cat. No. A9789) at 10 mg/mL.

Cerebral cortices were rapidly dissected in ice-cold buffer containing actinomycin D (5 µg/mL), triptolide (10 µM), and anisomycin (27.1 µg/mL). Cortices were enzymatically dissociated using a papain-based digestion protocol as previously described, with the same concentrations of actinomycin D, triptolide, and anisomycin added immediately before tissue incubation. Enzyme mixtures were not pre-incubated with inhibitors. After dissociation, debris was removed, and single-cell suspensions were filtered and prepared for single-cell sequencing.

### 2.2 scRNA-seq and scATAC-seq library preparation

Single-cell suspensions were processed using the 10x Genomics Chromium platform. scRNA-seq libraries were generated using Chromium Single Cell 3′ v3 chemistry, targeting approximately 10,000 cells per library. After cells had been allocated for scRNA-seq library preparation, the remaining suspension was used for nuclei isolation and scATAC-seq library preparation with Chromium Single Cell ATAC chemistry, targeting approximately 10,000 nuclei per library. Libraries were sequenced on an Illumina NovaSeq 6000 S4 platform, targeting approximately 500 million reads per library.

### 2.3 scRNA-seq and scATAC-seq data processing

Raw scRNA-seq data were processed using Cell Ranger v3.1.0 with the mm10-3.0.0 mouse transcriptome reference. Gene-barcode matrices were generated using the standard Cell Ranger pipeline with default parameters.

Raw scATAC-seq data were processed using Cell Ranger ATAC v2.2.0 with the 10x Genomics mouse reference refdata-cellranger-arc-mm10-2020-A-2.0.0. Fragment files, peak-barcode matrices, and quality-control outputs were generated using the standard Cell Ranger ATAC pipeline with default parameters. Sample integration and downstream analyses were performed in Seurat as described below.

## 3. Single-cell sequencing analysis

### 3.1 scRNA-seq analysis

Preprocessing followed^29^, with modifications described above. Full analysis code is available at https://github.com/thomaskim-lab/STAT1-Paper.

Microglia identified in each genotype-annotated parent dataset (WT, *Stat1* KO, and *Irf1* KO) were subset and merged pairwise for genotype-level comparisons (WT + *Stat1* KO and WT + *Irf1* KO). After merging, data were re-normalized using SCTransform, with nCount_RNA and nFeature_RNA regressed out, followed by principal component analysis (PCA). Harmony was used to integrate samples (group.by.vars = “SampleID”), and the corrected embedding was used for UMAP visualization and graph-based Louvain clustering.

An initial clustering pass was used to identify and remove low-quality or non-microglial contaminating clusters. The remaining cells were then re-normalized, re-integrated with Harmony, and re-clustered through one or two additional iterative rounds until a stable microglia-only cluster structure was obtained. Expression of interferon-response genes (*Ifit1, Ifit3, Isg15,* and *Irf7*) was visualized using feature plots to aid identification of interferon-response microglia (IRM). Final clusters were manually annotated as homeostatic, disease-associated microglia (DAM)-like, or IRM subtypes based on marker expression.

Cell-level marker-expression profiles across biological conditions (WT versus *Stat1* KO and WT versus *Irf1* KO) were evaluated using the final annotated microglia objects, with genotype condition specified as the grouping identity. For direct comparison of *Stat1* KO and *Irf1* KO states, single-cell profiles from each knockout condition were subset from their respective final objects and concatenated. Data were re-normalized using SCTransform while accounting for sequencing depth and feature counts through nCount_RNA and nFeature_RNA, respectively, and re-projected using PCA and UMAP. Condition-associated transcripts distinguishing *Stat1* KO from *Irf1* KO microglia were identified using FindMarkers.

#### 3.1.1 STAT1-APOE cross-referencing in public human and cross-species datasets

The association between STAT1 and APOE was examined in two publicly available, previously annotated single-cell or single-nucleus datasets^30,31^.

Cells were classified according to whether STAT1 transcript was detected in the log-normalized expression matrix, defined as values >0 versus values equal to 0. This detection-based classification was not intended as a curated biological positivity threshold and remains sensitive to transcript dropout, sequencing depth, and library complexity.

#### 3.1.2 Human AD dataset

Analyses were performed using a publicly available, annotated human microglial single-nucleus RNA-seq dataset^30,31^. Microglia had previously been classified into 12 transcriptional states, including homeostatic, lipid-processing, Inflammatory I, Inflammatory II, antiviral, and disease-associated intermediate populations. Donors had been staged as non-AD, early AD, or late AD.

The relationship between STAT1 detection status and APOE expression was tested using linear mixed-effects models implemented in lme4 and lmerTest: APOE expression ∼ STAT1 detection + log(1 + nCount) + nFeature + (1 | donor).

Models were fitted separately for each AD stage after pooling all microglial states and separately within each of the 12 transcriptional states. State-stage combinations containing fewer than 50 cells in total or fewer than 20 cells with detectable STAT1 transcript were excluded.

Gene specificity of the STAT1-APOE relationship was further examined in two individual microglial states. Within an inflammatory state, the same per-gene mixed-model framework was applied to *APOE, LIPA, LRRK2, FOXO3, FOXP1, SPON1, CCL3,* and *IL1B*. Within the lipid-processing state, the framework was applied to *APOE, LIPA, PPARG, ABCA1, TREM2,* and *LPL*.

#### 3.1.3 Cross-species disease-context dataset

A publicly available cross-species human/mouse microglial dataset integrated across disease contexts using Seurat reciprocal PCA (rPCA) was used to examine STAT1 and APOE co-expression across neurological disease states^29–31^.

Mouse conditions included experimental autoimmune encephalomyelitis, viral encephalitis, middle cerebral artery occlusion, familial AD models, graft-versus-host disease, and LPS challenge. Human conditions included AD, MS, amyotrophic lateral sclerosis, Parkinson’s disease, Huntington’s disease, glioma, and other neurological conditions represented in the source dataset.

Cells were assigned to four transcript-detection categories: STAT1 detected/APOE detected, STAT1 detected/APOE undetected, STAT1 undetected/APOE detected, or both transcripts undetected. The proportion of cells in each category was quantified by disease context or stage and visualized using stacked bar and pie charts separately for the human and mouse compartments.

Within each species, STAT1 expression was compared between cells with and without detectable *LPL* transcript using the Wilcoxon rank-sum test.

### 3.2 scATAC-seq preprocessing and per-sample processing

Each scATAC-seq sample was processed independently using Signac and Seurat. Fragment files were linked to the corresponding peak-by-cell objects, and gene annotations from EnsDb.Mmusculus.v79 were added using the mm10 genome assembly.

Each sample was independently normalized using term frequency-inverse document frequency (TF-IDF) normalization. Singular value decomposition was performed on the top features using min.cutoff = “q0” to generate a latent semantic indexing (LSI) embedding. UMAP visualization and graph-based Louvain clustering with multilevel refinement (algorithm = 3) were performed using LSI components 2-30.

Cell-type identities were assigned using per-cell gene-activity scores calculated with GeneActivity, followed by log-normalization of the activity matrix. Clusters were manually annotated based on the resulting markers, and microglia were subset from each sample for downstream analyses.

Detailed analysis parameters are provided in the GitHub repository cited above.

#### 3.2.1 Direct Stat1 knockout versus Irf1 knockout comparison

Microglia from *Stat1* KO and *Irf1* KO mice were merged and integrated using Harmony, with sample specified as the grouping variable and integration applied to the LSI embedding. The combined object was reprocessed using TF-IDF normalization, singular value decomposition, and UMAP based on LSI components 2-30.

Differential accessibility between *Stat1* KO and *Irf1* KO microglia was tested using logistic regression in FindMarkers (test.use = “LR”), with total peak counts included as a latent variable (latent.vars = “nCount_peaks”) to control for per-cell sequencing depth. Peaks were annotated to their nearest genes using ClosestFeature.

#### 3.2.2 Target-locus regulatory architecture and comparative analyses

##### 3.2.2.1 Building microglial enhancer annotations

Published H3K27ac and H3K4me1 ChIP-seq peak calls from microglia were obtained from the Ciernia laboratory reprocessed mm10 microglia epigenome track hub (github.com/ciernialab/MicrogliaChIPseq; TrackHubmm10_MGEpi_Master.txt). The hub aggregates reprocessed tracks from multiple published mouse microglial ChIP-seq studies, including^39,42–44^.

All bigBed tracks in the hub with filenames matching H3K27ac or H3K4me1 were downloaded and converted to BED format using bigBedToBed from the UCSC tools. Tracks were not restricted to a manually selected subset of contributing studies; the resulting collection therefore included all matching tracks available in the hub at the time of download.

BED files from all available H3K27ac tracks were concatenated, restricted to canonical chromosomes, sorted, and merged using bedtools merge to generate a combined microglial H3K27ac annotation. H3K4me1 tracks were processed in the same manner where available.

A permissive enhancer set was defined as merged H3K27ac intervals after subtraction of all genome-wide 2-kb promoter regions using bedtools subtract. The resulting intervals were re-merged and computationally validated to confirm the absence of residual promoter overlaps.

A strict enhancer set was defined as merged H3K27ac intervals directly overlapping merged H3K4me1 intervals using bedtools intersect -u. Genome-wide promoters were subsequently subtracted, and the absence of residual promoter overlaps was confirmed computationally.

Both enhancer sets were used exclusively as external annotations for the enrichment analyses described below and were not interpreted as direct evidence of STAT1 or IRF1 binding.

##### 3.2.2.2 Peak universe and genome-wide promoter annotation

The consensus peak set and associated count matrix were exported from the processed microglial object. Genome-wide gene annotations were used to define promoters as transcription start sites (TSS) ±2 kb for the primary analysis and TSS ±1 kb for sensitivity analyses. Promoter intervals were merged into non-overlapping genome-wide masks for downstream exclusion of promoter-overlapping distal regions.

##### 3.2.2.3 Target-gene compartment definitions

For each target lipid-handling gene, genomic compartments were defined as follows: promoters, TSS ±2 kb, with TSS ±1 kb used as a sensitivity definition; whole loci, TSS ±100 kb; and distal compartments, TSS ±100 kb after subtraction of all genome-wide 2-kb promoter regions.

An alternative whole-locus definition comprising the gene body ±100 kb was also generated. Exclusion of promoters from distal compartments was computationally validated to confirm the absence of residual overlap with the genome-wide promoter mask. Peaks were assigned to compartments based on genomic overlap, and peak-to-compartment maps and gene-level peak counts were tabulated.

##### 3.2.2.4 Promoter, distal, and locus differential-accessibility summarization

Pseudobulk differential-accessibility results for *Stat1* KO versus WT and *Irf1* KO versus WT were joined to the target-gene compartment map.

For each gene, genotype comparison, and genomic compartment, the mean and median log fold changes, numbers of gained and lost peaks at two false-discovery-rate (FDR) thresholds, and mean log counts per million were summarized.

A redistribution score was calculated for each gene as the distal mean log fold change minus the promoter mean log fold change, summarizing the relative redistribution of accessibility from promoter to distal regions.

##### 3.2.2.5 Matched-control gene analysis

To assess whether target lipid-handling genes showed distinct accessibility redistribution relative to the broader genome, non-target control genes were selected by nearest-neighbor matching in a scaled feature space.

Matching variables included log1p-transformed baseline WT promoter accessibility, baseline WT whole-locus accessibility, baseline WT distal accessibility, gene length, and peak counts within each compartment. Features with zero variance were removed before scaling. Euclidean distance was used to identify 20 matched control genes for each target gene.

One-sided Wilcoxon rank-sum tests compared target genes with their matched controls for promoter log fold change, hypothesized to be lower in target genes; distal log fold change and redistribution score, hypothesized to be higher; and the numbers of significantly altered peaks within each compartment. *P* values were adjusted using the Benjamini-Hochberg procedure within each genotype comparison and target-gene-class grouping.

##### 3.2.2.6 Enhancer annotation enrichment

Distal peaks altered in each genotype comparison were tested for overlap with published microglial enhancer annotations. The permissive annotation comprised promoter-excluded H3K27ac-marked regions, whereas the strict annotation comprised promoter-excluded regions co-marked by H3K27ac and H3K4me1.

For each query peak set, defined by genotype, direction of change, significance threshold, and, where applicable, restriction to lipid-associated loci, a matched background was constructed from the complete distal peak universe.

Background peaks were matched by binning baseline WT accessibility into 20 bins, peak width into 10 bins, and distance to the nearest TSS into 20 bins. Chromosome and bin-matching criteria were progressively relaxed when insufficient candidates were available, and approximately 20 background peaks were sampled per query peak.

One-sided Fisher’s exact tests, with the alternative hypothesis specified as greater, were used to assess enrichment of enhancer overlap among query peaks relative to matched-background peaks. *P* values were adjusted using the Benjamini-Hochberg procedure within each enhancer-annotation set.

A parallel analysis assessed enrichment among distal peaks altered in the same direction in both *Stat1* KO and *Irf1* KO microglia across nominal *P*-value and FDR thresholds.

##### 3.2.2.7 Shared and factor-biased candidate peak classification

To compare accessibility changes between *Stat1* KO and *Irf1* KO conditions across the peak universe, differential-accessibility metrics were merged at the peak level. Regions were assigned to relative-response categories, including shared concordant, discordant, and conditionally enriched groups, according to directional agreement and statistical confidence across the two comparisons.

To avoid attributing apparent factor exclusivity to threshold-boundary effects or differences in statistical power between cohorts, these categories were treated solely as descriptive groupings of relative response magnitude and not as evidence of unique transcription-factor recruitment or direct mechanistic exclusivity.

## 4. HMC3 cell culture

HMC3 human microglial cells were obtained directly from ATCC (CRL-3304) and used between passages 5 and 20. Cells were maintained in DMEM (Cat. No. 41966029) supplemented with 10% heat-inactivated fetal bovine serum (FBS; Cat. No. SH30071.03HI) and 1× penicillin-streptomycin (Cat. No. 15140122).

Cells were cultured at 37°C in a humidified incubator with 5% CO. Medium was replaced every 3 days, and cells were passaged at 70-80% confluency using TrypLE (Cat. No. 12604013) after washing with PBS. Detached cells were collected by centrifugation at 300 × g for 5 min and resuspended in complete medium for subsequent plating.

### 4.1 LPS treatment of HMC3 cells

For inflammatory stimulation, HMC3 cells were treated with lipopolysaccharide (LPS; Cat. No. 15536286) at 5 µg/mL in complete medium under the same serum conditions as untreated controls.

LPS-containing medium was prepared immediately before use and added after removal of the original culture medium. Control cells received complete medium without LPS. Cells were treated for 24 or 48 h before RNA extraction, qPCR, SABER-FISH, or bulk RNA-seq analysis, as specified for each experiment.

### 4.2 STAT1 siRNA knockdown in HMC3 cells

STAT1 knockdown was performed using a TriFECTa Dicer-substrate RNAi kit targeting human STAT1 (IDT, hs.Ri.STAT1.13, Cat. No. 514774154) and Lipofectamine RNAiMAX transfection reagent (Cat. No. 13778100). A non-targeting DsiRNA was used as the control.

DsiRNAs were resuspended according to the manufacturer’s instructions and diluted in Opti-MEM I Reduced Serum Medium (Cat. No. 31985062) before transfection. A final DsiRNA concentration of 10 nM was used in all knockdown experiments.

For transfection in 12-well plates, complexes were prepared in Opti-MEM using 1.5 µL Lipofectamine RNAiMAX and 5 µL of a 1 µM DsiRNA working stock per well. Complexes were incubated for 10-20 min at room temperature before addition to cells in antibiotic-free complete medium.

After transfection, cells were returned to the incubator without a medium change unless replacement was required because of cell condition. For experiments combining STAT1 knockdown with inflammatory stimulation, LPS treatment was initiated after siRNA transfection at the time specified for each experiment. Cells were harvested at the indicated time points for qPCR, SABER-FISH, flow cytometry, or bulk RNA-seq analysis.

### 4.3 Plasmid-based STAT1 overexpression rescue

To assess whether restoration of STAT1 expression could rescue the phenotype observed after siRNA knockdown, HMC3 cells were transfected with an N-terminal eGFP-tagged human STAT1 expression plasmid (eGFP STAT1 WT; Addgene plasmid #12301) carrying a codon-optimized STAT1 coding sequence after the STAT1 knockdown procedure described in Section 4.2.

Plasmid transfection was performed using Lipofectamine 3000 Transfection Reagent (Thermo Fisher Scientific, Cat. No. L3000015) according to the manufacturer’s protocol. Briefly, two solutions were prepared separately: Lipofectamine 3000 reagent diluted in Opti-MEM I Reduced Serum Medium and plasmid DNA plus P3000 reagent diluted in Opti-MEM. The solutions were combined, and transfection complexes were incubated for 15-20 min at room temperature.

For each well, 100 µL of transfection complex was added directly to the cells, followed by 900 µL of pre-warmed complete growth medium consisting of DMEM supplemented with 10% FBS without antibiotics. Cells were incubated with the transfection mixture for 4 h, after which the medium was replaced with fresh, pre-warmed complete medium. Cells were harvested at the indicated time points for downstream functional and transcriptional analyses.

### 4.4 RNA extraction, cDNA synthesis, and quantitative PCR

Total RNA was isolated from HMC3 cells using the ReliaPrep RNA MiniPrep Cells kit (Cat. No. Z6014) or Monarch Spin RNA Isolation Kit (Cat. No. NEB-T2110S), according to the manufacturers’ instructions.

First-strand cDNA was synthesized using SuperScript IV Reverse Transcriptase (Cat. No. 18090010) according to the manufacturer’s instructions.

Quantitative PCR was performed using GoTaq qPCR Master Mix (Cat. No. A6001) or PowerUp SYBR Green Master Mix (Cat. No. A25742) on a StepOne Real-Time PCR System (Applied Biosystems). qPCR primers were obtained from IDT and designed using PrimerBank^45^.

Relative transcript abundance was quantified using the ΔΔCt method after normalization to the indicated internal reference genes. Primer sequences and reference genes used for each experiment are provided in Table S20.

## 5. Bulk RNA sequencing

### 5.1 HMC3 LPS experiments

For HMC3 LPS bulk RNA-seq experiments, polyadenylated mRNA was reverse-transcribed using barcoded primers containing unique molecular identifiers and template-switching sequences to generate full-length cDNA, which was subsequently amplified.

Amplified cDNA was processed using an Illumina-compatible library-preparation workflow with a 10x Genomics library-construction kit. Final libraries underwent quality control and were sequenced on an Illumina NovaSeq X Plus platform.

Raw FASTQ files were aligned to the *Homo sapiens* GRCh38 reference genome using Ensembl release 108 annotations and STAR v2.7.11b. Reads were mapped with unique mapping enabled, and unmapped reads were retained. UMI sequences were extracted and deduplicated.

Alignment statistics and gene-level count matrices were generated for downstream analysis.

### 5.2 Plasmidsaurus RNA-seq

For additional bulk transcriptomic profiling, cell pellets were submitted to Plasmidsaurus for 3′ digital gene-expression (DGE) RNA-seq. Cells were harvested by scraping or TrypLE detachment, pelleted by centrifugation, and resuspended in 1× DNA/RNA Shield (Cat. No. R1100).

Lysates were homogenized by pipetting and submitted at ambient temperature according to the provider’s sample-submission guidelines. Mouse samples were submitted as *Mus musculus* and human samples as *Homo sapiens*.

Libraries were prepared and sequenced by Plasmidsaurus using its DGE RNA-seq workflow. The provider returned approximately 10 million deduplicated Illumina reads per sample together with gene-level count tables, deduplicated read outputs, and sequencing quality-control metrics. These outputs were used for downstream analyses.

### 5.3 Bulk RNA-seq analysis

Bulk RNA-seq analyses were performed in R using custom scripts. Gene-level count matrices and sample-annotation files were imported from provider-generated outputs or STAR-derived count tables, depending on the dataset. For Plasmidsaurus DGE RNA-seq datasets, expression matrices containing gene-level count and counts-per-million values were imported, and raw count columns were extracted for differential-expression analysis. For the HMC3 LPS bulk RNA-seq dataset, STAR-generated raw count matrices were imported together with the corresponding sample metadata.

Ensembl gene identifiers were stripped of version suffixes where present. Duplicated Ensembl identifiers were removed, and count matrices were rounded and converted to integer mode before DESeq2 analysis. Sample metadata were parsed from sample names and manually verified against sample-annotation tables. Depending on the dataset, metadata fields included cell type, species, treatment, time point, knockdown condition, siRNA duplex, replicate identity, sequencing order or batch, and combined experimental group. Dataset- and sample-level information, including Plasmidsaurus order identifiers, species, experimental conditions, replicate numbers, and primary contrasts, is provided at https://github.com/thomaskim-lab/STAT1-Paper.

Differential-expression analysis was performed using DESeq2. Before model fitting, low-count genes were filtered by retaining genes with counts ≥10 in at least two samples. For the HMC3 LPS time-course analysis, DESeq2 models included time point, treatment, and their interaction. LPS-treated samples were compared with matched controls at each time point. For HMC3 STAT1-knockdown experiments without LPS, independent Plasmidsaurus sequencing orders were merged and analyzed using a batch-adjusted model with the design: ∼ Batch + Condition. Individual STAT1 DsiRNA contrasts and pooled STAT1-knockdown contrasts were tested against control-transfected cells. For HMC3 STAT1 knockdown combined with LPS treatment, both factorial and group-based models were used. The factorial model used the design: ∼ Knockdown + LPS + Knockdown. This model estimated the main effects of STAT1 knockdown and LPS treatment and their interaction. Group-based models were used for direct pairwise comparisons among control, STAT1 knockdown, LPS-treated control, and STAT1 knockdown plus LPS conditions.

For primary mouse microglia *Stat1*-knockdown experiments, untreated samples were retained for baseline quality control but excluded from the primary differential-expression contrasts. Lipofectamine-only controls were compared with individual *Stat1* DsiRNA conditions and pooled *Stat1*-knockdown samples.

Gene annotations were added using AnnotationDbi with org.Hs.eg.db for human datasets and org.Mm.eg.db for mouse datasets. Differential-expression result tables were exported with Ensembl identifiers, gene symbols, Entrez identifiers, gene names, base mean expression, log2 fold change, standard error, test statistic, nominal *P* value, and Benjamini-Hochberg-adjusted *P* value. Genes were considered differentially expressed at an adjusted *P*-value threshold of 0.05. Additional log2 fold-change thresholds were applied as specified for visualization, top-gene selection, and pathway analysis. Variance-stabilizing transformation was performed using DESeq2 with blind = FALSE. Transformed expression values were used for PCA, sample-distance heatmaps, selected-gene heatmaps, and module-score calculations. Sample distances were calculated from variance-stabilized expression matrices using Euclidean distance. Heatmaps were generated from variance-stabilized expression values, with gene-wise scaling where indicated. Gene Ontology biological-process enrichment was performed using clusterProfiler on significantly upregulated or downregulated genes, using Entrez identifiers and Benjamini-Hochberg correction. Reactome pathway enrichment was assessed using ReactomePA.

For ranked pathway analyses, genes were ranked primarily according to the DESeq2 Wald statistic. When the

Wald statistic was unavailable, genes were ranked using the signed −log_10_ *P* value multiplied by the direction of the log2 fold change. Reactome gene-set enrichment analysis was performed using these ranked gene lists. Hallmark gene-set enrichment analysis was performed for human datasets, where indicated, using MSigDB Hallmark gene sets obtained through msigdbr and analyzed with fgsea. Focused analyses used curated gene modules related to lipid-droplet structure, lipolysis, lipid esterification, fatty-acid activation and oxidation, lysosome and lipophagy, endoplasmic-reticulum stress, interferon, STAT and IRF signaling, NF-κB-associated inflammation, microglial identity, disease-associated microglia, antigen presentation, oxidative phosphorylation, glycolysis, and iron, redox, and glutathione biology.

For each contrast, module-level summaries were calculated from the differential-expression results as the mean and median log_2_ fold change among detected module genes and the numbers of significantly upregulated and downregulated genes within each module. For selected datasets, sample-level module scores were calculated from variance-stabilized expression matrices after collapsing Ensembl identifiers to gene symbols by retaining the highest-expressed Ensembl identifier for each symbol. Each gene was z-scored across samples, and module scores were calculated as the mean gene-wise z-score across genes represented in the module. Module-score differences were tested using linear models incorporating knockdown, LPS treatment, and their interaction where applicable.

## 6. Primary mouse microglia culture

Primary microglia were prepared as previously described^46^, with minor modifications. Briefly, brains from postnatal day 1-2 pups were collected in ice-cold L15 medium, mechanically dissociated, and plated as mixed glial cultures in polyethylenimine-coated flasks containing low-glucose DMEM supplemented with 10% heat-inactivated FBS and 1% penicillin-streptomycin.

Cultures were maintained at 37°C in a humidified incubator with 5% CO. At 48 h after plating, 8 mL of the 10-mL culture volume was replaced with fresh low-glucose DMEM containing 10% heat-inactivated FBS and 1% penicillin-streptomycin. Mixed glial cultures were maintained for 10-14 days to allow progressive microglial accumulation and were subsequently supplemented every 24 h with 1 mL low-glucose DMEM containing 1% heat-inactivated FBS and 1% penicillin-streptomycin. Microglia were used within approximately 10 days after purification.

For experimental plating, purified microglia were enzymatically detached using TrypLE Express, collected by centrifugation at 1,300 × g for 6 min at 4°C, and resuspended in low-glucose DMEM containing 1% heat-inactivated FBS, 1% penicillin-streptomycin, and recombinant mouse M-CSF (PeproTech, Cat. No. 315-02) at 10 ng/mL. Viable cells were counted by trypan blue exclusion and plated at 110,000 cells/cm². Cells were allowed to adhere and recover overnight before experimental treatment or transfection.

### 6.1 STAT1 siRNA knockdown in primary mouse microglia

STAT1 knockdown in primary mouse microglia was performed using a TriFECTa Dicer-substrate RNAi kit targeting mouse *Stat1* (IDT, mm.Ri.Stat1.13, Cat. No. 6746061) and Lipofectamine RNAiMAX, following the general transfection workflow described for HMC3 cells. A non-targeting DsiRNA was used as the control.

DsiRNA was used at a final concentration of 10 nM. After transfection, cells were maintained in primary microglia culture medium supplemented with recombinant mouse M-CSF at 10 ng/mL.

Cells were harvested at the indicated time points for qPCR, flow cytometry, and/or DGE RNA-seq analysis. qPCR primer sequences are provided in Table S20.

### 6.2 Primary microglia rescue treatments and lipid-peroxidation assays

Following *Stat1* knockdown, primary mouse microglia were treated with recombinant mouse IFNγ, ferrostatin-1, the PPARγ agonist pioglitazone, or oleate-BSA, as specified for each experiment. Recombinant mouse IFNγ (Cat. No. 575302) was used at 50-100 ng/mL. Ferrostatin-1 (Cat. No. HY-100579) and pioglitazone (Cat. No. HY-13956) were prepared according to the manufacturers’ recommendations and used at 1-5 µM. Oleate-BSA was used at 20-250 µM. Treatments were applied for 18-24 h before harvest, with vehicle-matched controls included for each treatment.

Lipid peroxidation was measured using the BODIPY 581/591 C11 lipid-peroxidation sensor (Cat. No. D3861). After the indicated treatments, culture medium was replaced with serum-free medium containing BODIPY 581/591 C11 at 1:1,000 from a 1 mg/mL stock solution. Cells were incubated for 30 min at 37°C before harvest and acquisition by flow cytometry as described below.

## 7. Flow Cytometry

Following the indicated treatments, cells were washed with PBS, detached with TrypLE for 5 min, and collected by centrifugation at 500 × g for 5 min at 4°C. Cells were then stained with the indicated lipid dyes, viability dyes, or antibodies and resuspended in flow-cytometry buffer consisting of PBS containing 0.5% BSA and 2 mM EDTA.

Neutral-lipid-associated fluorescence was assessed using BODIPY 493/503 (Cat. No. D3922) or HCS LipidTOX Red Neutral Lipid Stain (Cat. No. H34476), as specified for each experiment. Cells were stained at 1:1,000 from a 1 mg/mL stock solution in PBS for 10 min at 37°C followed by 10 min at room temperature. Where applicable, Zombie NIR was then added at 1:1,000 in PBS according to the manufacturer’s instructions. Cells were subsequently resuspended in flow-cytometry buffer, and DAPI was added at 1 µg/mL immediately before acquisition where applicable.

Samples were acquired using a NovoCyte Quanteon flow cytometer (Agilent Technologies, Inc., Santa Clara, CA, USA). Forward- and side-scatter parameters were used to exclude debris, followed by doublet exclusion and live-cell gating where applicable. Median fluorescence intensities of neutral lipid probes were quantified using FlowJo v10. Representative gating strategies are shown in Supplementary Figs. S7C and S10E.

## 8. Single-molecule fluorescent *in situ* hybridization (smfISH)

SABER-FISH was performed in cultured cells based on previously described methods^47^, with minor modifications. Probes targeting human and mouse transcripts were designed using OligoMiner-derived probe resources and custom filtering workflows against hg38 or mm10 genomic annotations.

Candidate exon-targeting probes were filtered for sequence redundancy and melting temperature, and 24-50 probes were selected per target transcript. Probe oligonucleotides were appended with a TTT linker and a 9-mer primer-exchange-reaction (PER) primer sequence. Probe identities, target genes, PER barcodes, and fluorophores are listed in Table S20.

Probe oligonucleotides were pooled by target gene and extended by PER using Bst large-fragment polymerase (Cat. No. BPL-300). PER products were purified using Monarch PCR and DNA Cleanup columns, quantified using a NanoDrop spectrophotometer, and used for hybridization.

Cells were cultured in ibidi 8-well high-polymer chambers with a #1.5 polymer coverslip and ibiTreat surface treatment (Cat. No. 80806). Fixed cells were permeabilized, equilibrated in SSC-based hybridization buffer, and hybridized with pooled PER-concatemerized probe sets at 42°C.

After primary hybridization and washing, fluorescent oligonucleotides complementary to the PER barcodes were hybridized at 37°C. For multiplexed imaging, fluorescent signals were stripped using a formamide-containing displacement buffer, and samples were re-hybridized with subsequent fluorescent oligonucleotide sets over iterative imaging rounds.

Images were acquired using an Olympus APEXVIEW APX100 fluorescence microscope (Evident Corporation) equipped with a Hamamatsu camera, motorized multi-position acquisition, and fluorescence-filter sets appropriate for the fluorophores used in each experiment. Images were acquired with a 40× objective using tiled XY acquisition and z-stacks. Identical imaging regions were re-imaged across sequential hybridization rounds using the microscope well-navigation function.

Cell segmentation was performed using ilastik^48^, and SABER-FISH spot detection and quantification were performed using RS-FISH^49^. Spot counts were quantified within segmented cellular regions.

## 9. Immunocytochemistry and imaging

Cells were seeded in µ-Plate 24 Well Black plates with an ibiTreat #1.5 polymer coverslip bottom (ibidi GmbH). After the indicated treatments, cells were fixed with 4% PFA for 20 min at room temperature, washed with 1× PBS, and stained with HCS LipidTOX Red Neutral Lipid Stain at 1:1,000 for 30 min at 37°C. Nuclei were stained with DAPI at 1 µg/mL.

Imaging was performed at 40× magnification using an Olympus APX100 automated fluorescence microscope equipped with a DP23M-CU monochrome CMOS camera and operated using cellSens software.

Quantification was performed in FIJI/ImageJ v2.16.0 using manually segmented cells for each condition. Background fluorescence was determined from cell-free regions and subtracted from cellular measurements. Integrated density was then divided by cell area to obtain fluorescence intensity per unit area, thereby accounting for differences in cell size.

## Data availability

Sequencing data generated in this study are being deposited in the ArrayExpress collection at EMBL-EBI BioStudies. Some sequencing data are available in the Gene Expression Omnibus under accession GSE175546.

## Code availability

Custom computer code and analysis scripts used in this study are available at https://github.com/thomaskim-lab/STAT1-Paper.

## Author contributions

A.E.M. and D.W.K. conceived the study. A.E.M., J.M.B.L., N.D.J., Y.L. and L.L. performed experiments. A.E.M. and D.W.K. analyzed the data. D.W.K. supervised the study, acquired funding and administered the project. A.E.M. and D.W.K. wrote the manuscript.

## Acknowledgements

Flow cytometry was performed at the FACS Core Facility, Aarhus University, Denmark. RNA sequencing was performed at the Stenomics Core Facility, Aarhus University, Denmark. Some of the computational work for this project was performed on the GenomeDK cluster. We thank GenomeDK and Aarhus University for providing computational resources and support that contributed to this research.

## Supplemental Table Legends

**Table S1.** Differentially expressed genes in *Stat1*-/- versus WT astrocytes.

**Table S2.** Differentially expressed genes in *Stat1*-/- versus WT endothelial cells.

**Table S3.** Differentially expressed genes in *Stat1*-/- versus WT neurons.

**Table S4**. Differentially expressed genes in *Stat1*-/- versus WT microglia.

**Table S5.** Differentially expressed genes in *Irf1*-/- versus WT microglia.

**Table S6.** Differentially expressed genes in *Irf1*-/- versus WT astrocytes.

**Table S7.** Differentially expressed genes in *Irf1*-/- versus WT endothelial cells.

**Table S8.** Differentially expressed genes in *Irf1*-/- versus WT neurons.

**Table S9.** Differentially expressed genes in *Irf1*-/- versus *Stat1*-/- microglia.

**Table S10.** Candidate differential-accessibility peaks in WT, *Stat1*-/-, and *Irf1*-/- microglia, with shared and factor-biased response classifications.

**Table S11.** Differentially expressed genes in primary microglia following acute *Stat1* siRNA depletion.

**Table S12.** Projection scores of primary microglial bulk RNA-seq signatures onto the adult in vivo microglial landscape.

**Table S13.** Differentially expressed genes and module scores in primary microglia following acute IFNγ treatment.

**Table S14.** Differentially expressed genes in control primary microglia following pioglitazone treatment.

**Table S15.** Differentially expressed genes in *Stat1*-depleted primary microglia following pioglitazone treatment.

**Table S16.** Differentially expressed genes in HMC3 cells following 24 h LPS treatment.

**Table S17.** Differentially expressed genes in HMC3 cells following 48 h LPS treatment.

**Table S18.** Differentially expressed genes in HMC3 cells following 24 h STAT1 siRNA depletion.

**Table S19.** Differentially expressed genes in HMC3 cells following 48 h STAT1 siRNA depletion.

**Table S20.** Quantitative PCR primer sequences and SABER-FISH probe set specifications.

---

**Figure S1.**
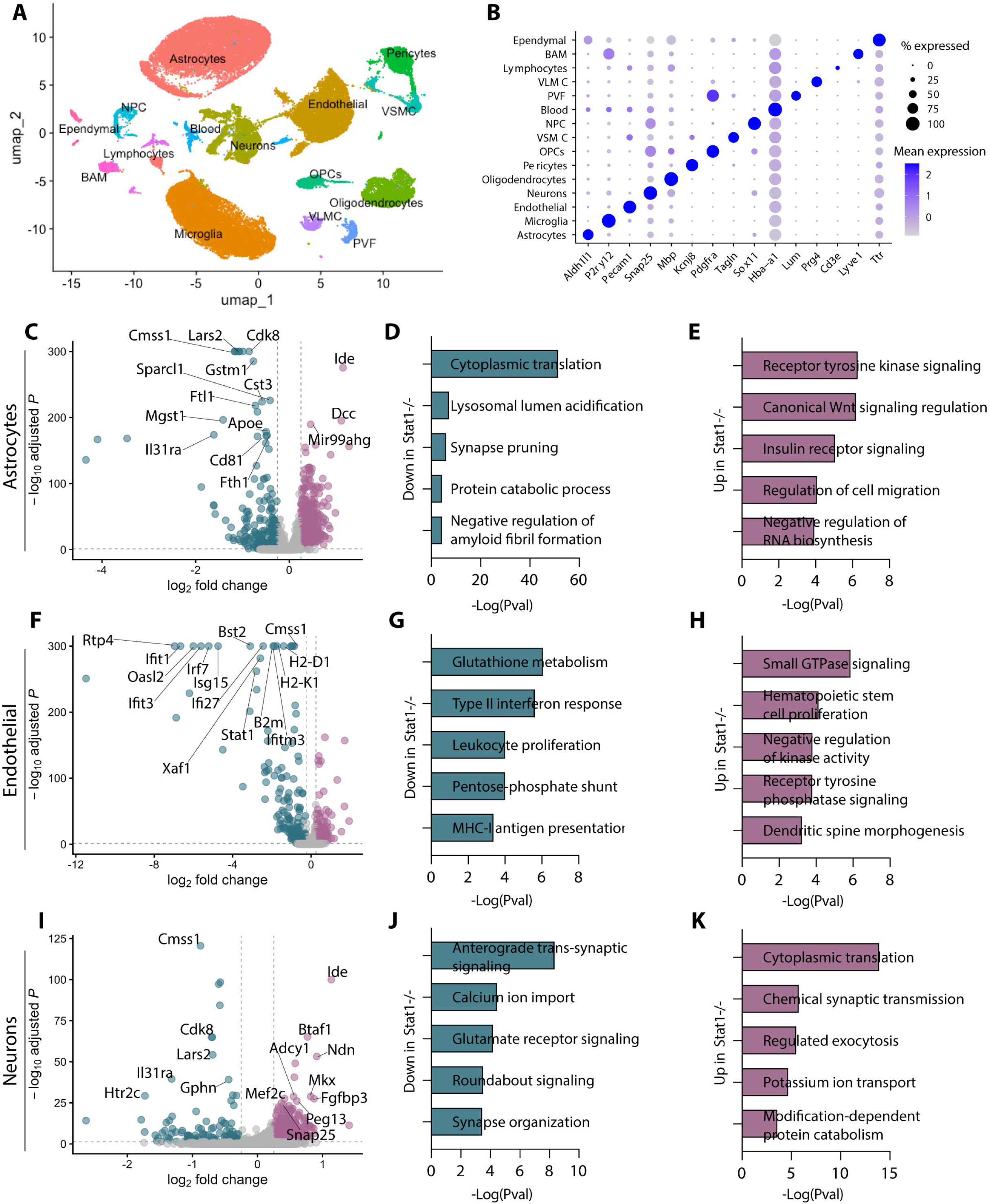
STAT1 deficiency produces cell-type-specific transcriptional changes across the adult cerebral cortex. **(A)** UMAP visualization of the complete adult mouse brain scRNA-seq dataset, showing the major neural, glial, vascular, and immune cell populations identified by transcriptional clustering and canonical lineage markers. BAM, border-associated macrophages; NPC, neural progenitor cells; OPC, oligodendrocyte precursor cells; PVF, perivascular fibroblasts; VLMC, vascular and leptomeningeal cells; VSMC, vascular smooth muscle cells. **(B)** Dot plot of representative canonical marker genes across the annotated cell populations. Dot size indicates the percentage of cells expressing each gene, and color indicates scaled mean expression. **(C)** Volcano plot of differential gene expression between *Stat1*−/− and WT astrocytes. **(D, E)** Selected biological process terms enriched among genes expressed at lower (D) or higher (E) levels in *Stat1*−/− relative to WT astrocytes. **(F)** Volcano plot of differential gene expression between *Stat1*−/− and WT endothelial cells. (G, H) Selected biological process terms enriched among genes expressed at lower (G) or higher (H) levels in *Stat1*−/− relative to WT endothelial cells. **(I)** Volcano plot of differential gene expression between *Stat1*−/− and WT neurons. **(J, K)** Selected biological process terms enriched among genes expressed at lower (J) or higher (K) levels in *Stat1*−/− relative to WT neurons. In (C), (F), and (I), genes are categorized according to whether they meet the fold-change threshold, the statistical-significance threshold, both, or neither; selected genes are labeled. In (D), (E), (G), (H), (J), and (K), bar length indicates −log_10_(P value).

**Figure S2.**
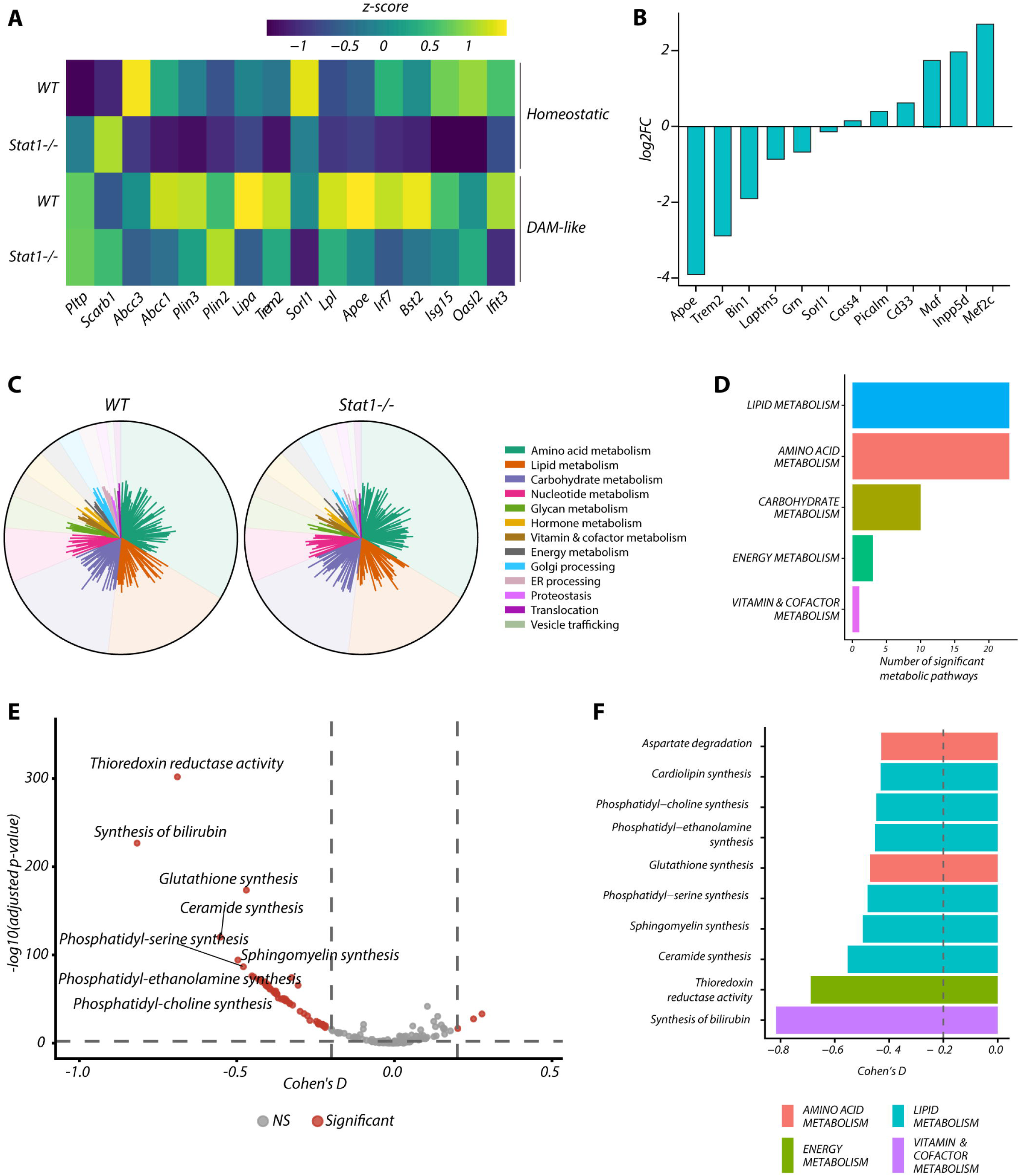
STAT1 loss alters state-resolved lipid-associated transcription and predicted metabolic programs in microglia. **(A)** Heatmap of scaled expression of selected lipid-handling, microglial-state, and interferon-response genes in homeostatic and DAM-like microglia from WT and *Stat1*−/− mice. Colors indicate row-scaled z-scores. **(B)** Log fold changes for selected Alzheimer’s disease risk-associated and microglial genes in *Stat1*−/− relative to WT microglia. Negative values indicate lower and positive values higher expression in *Stat1*−/− microglia. **(C)** Global comparison of transcriptome-derived metabolic-task profiles in WT and *Stat1*−/− microglia. Predicted tasks are grouped by functional category, including amino-acid, lipid, carbohydrate, nucleotide, glycan, hormone, vitamin and cofactor, and energy metabolism, together with cellular processing pathways. **(D)** Numbers of significantly altered predicted metabolic tasks in *Stat1*−/− relative to WT microglia, grouped by metabolic category. **(E)** Differential analysis of transcriptome-derived metabolic tasks between WT and *Stat1*−/− microglia. Each point represents a predicted metabolic task; the x-axis indicates Cohen’s d effect size and the y-axis −log₁₀ adjusted P value. Significantly altered tasks are highlighted and selected pathways labeled. Dashed lines indicate the significance and effect-size thresholds. Negative values indicate lower predicted task scores in *Stat1*−/− microglia. **(F)** Effect sizes for selected metabolic tasks significantly altered in *Stat1*−/− relative to WT microglia, including phospholipid, sphingolipid, ceramide, cardiolipin, glutathione, redox, and vitamin or cofactor-associated processes. Bars are colored by broad metabolic category, and the dashed vertical line indicates the effect-size threshold used for selection.

**Figure S3.**
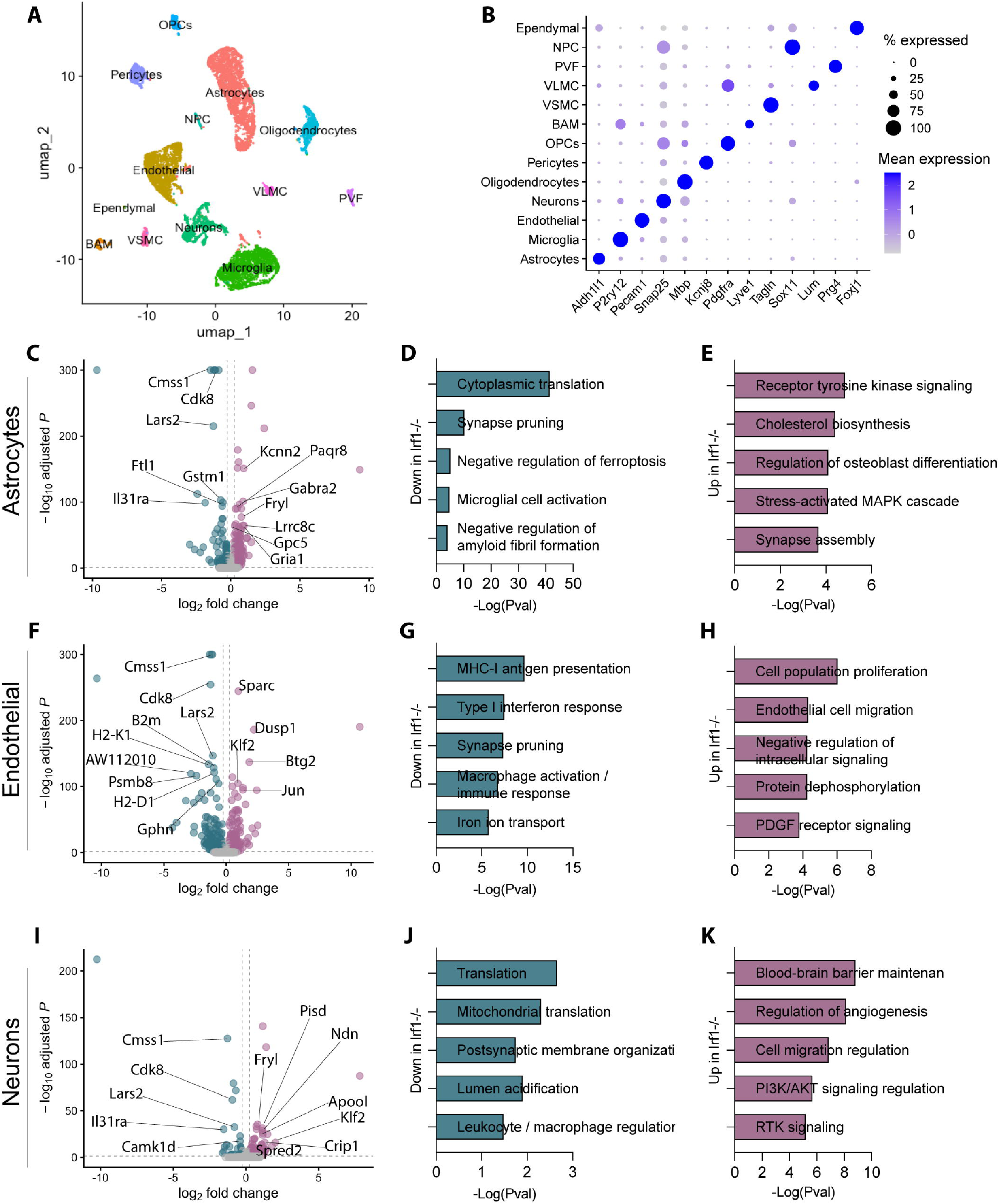
IRF1 deficiency produces cell-type-specific transcriptional changes across the adult cerebral cortex. **(A)** UMAP visualization of the complete adult mouse brain scRNA-seq dataset from WT and *Irf1*−/− mice, showing the major neural, glial, vascular, and immune cell populations identified by transcriptional clustering and canonical lineage markers. BAM, border-associated macrophages; NPC, neural progenitor cells; OPC, oligodendrocyte precursor cells; PVF, perivascular fibroblasts; VLMC, vascular and leptomeningeal cells; VSMC, vascular smooth muscle cells. **(B)** Dot plot of representative canonical lineage-marker genes across the annotated cell populations. Dot size indicates the percentage of cells expressing each gene, and color indicates scaled mean expression. **(C)** Volcano plot of differential gene expression between *Irf1*−/− and WT astrocytes. **(D, E)** Selected biological process terms enriched among genes expressed at lower (D) or higher (E) levels in *Irf1*−/− relative to WT astrocytes. **(F)** Volcano plot of differential gene expression between *Irf1*−/− and WT endothelial cells. **(G, H)** Selected biological process terms enriched among genes expressed at lower (G) or higher (H) levels in *Irf1*−/− relative to WT endothelial cells. **(I)** Volcano plot of differential gene expression between *Irf1*−/− and WT neurons. **(J, K)** Selected biological process terms enriched among genes expressed at lower (J) or higher (K) levels in *Irf1*−/− relative to WT neurons. In (C), (F), and (I), genes are categorized according to whether they meet the fold-change threshold, the statistical-significance threshold, both, or neither; selected genes are labeled. In (D), (E), (G), (H), (J), and (K), bar length indicates −log_10_(P value).

**Figure S4.**
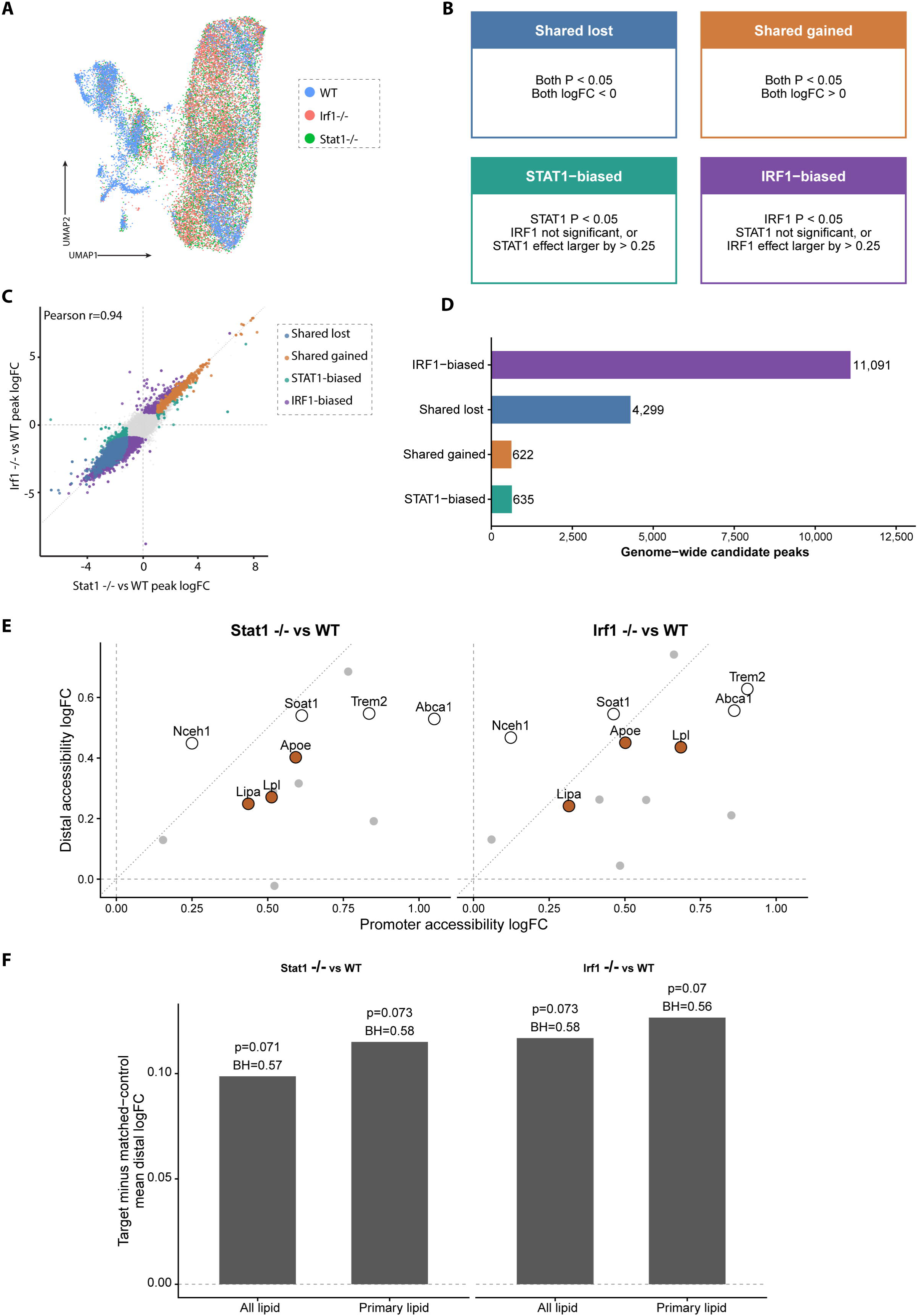
Loss of STAT1 or IRF1 produces concordant genome-wide chromatin-accessibility changes with shared and factor-biased responses at lipid-handling loci. **(A)** Integrated UMAP visualization of single-cell chromatin-accessibility profiles from WT, *Stat1*−/−, and *Irf1*−/− microglia, colored by genotype. **(B)** Classification scheme used to assign candidate differential-accessibility peaks to shared lost, shared gained, STAT1-biased, or IRF1-biased response classes. **(C)** Comparison of genome-wide candidate-peak accessibility effect sizes in *Stat1*−/− versus WT and *Irf1*−/− versus WT microglia. Candidate peaks are colored according to the response classes defined in (B). **(D)** Numbers of genome-wide candidate differential-accessibility peaks assigned to each response class. **(E)** Relationship between promoter and distal chromatin-accessibility changes at selected lipid-handling loci in *Stat1*−/− versus WT microglia (left) and *Irf1*−/− versus WT microglia (right). Each point represents a locus positioned according to its promoter and distal accessibility log fold changes. *Apoe*, *Lpl*, and *Lipa* are labeled in orange; open circles indicate additional genes discussed in the text, and grey points the remaining analyzed lipid-handling genes. Dashed horizontal and vertical lines indicate zero change, and the diagonal reference line indicates equal promoter and distal effects. **(F)** Distal-accessibility effects at lipid-handling target loci relative to matched control regions. Bars indicate the target-minus-matched-control mean distal accessibility log fold change for all lipid-handling loci and for the primary lipid-gene subset, in the *Stat1*−/− versus WT and *Irf1*−/− versus WT comparisons. Nominal and Benjamini-Hochberg-adjusted P values are shown above the bars.

**Figure S5.**
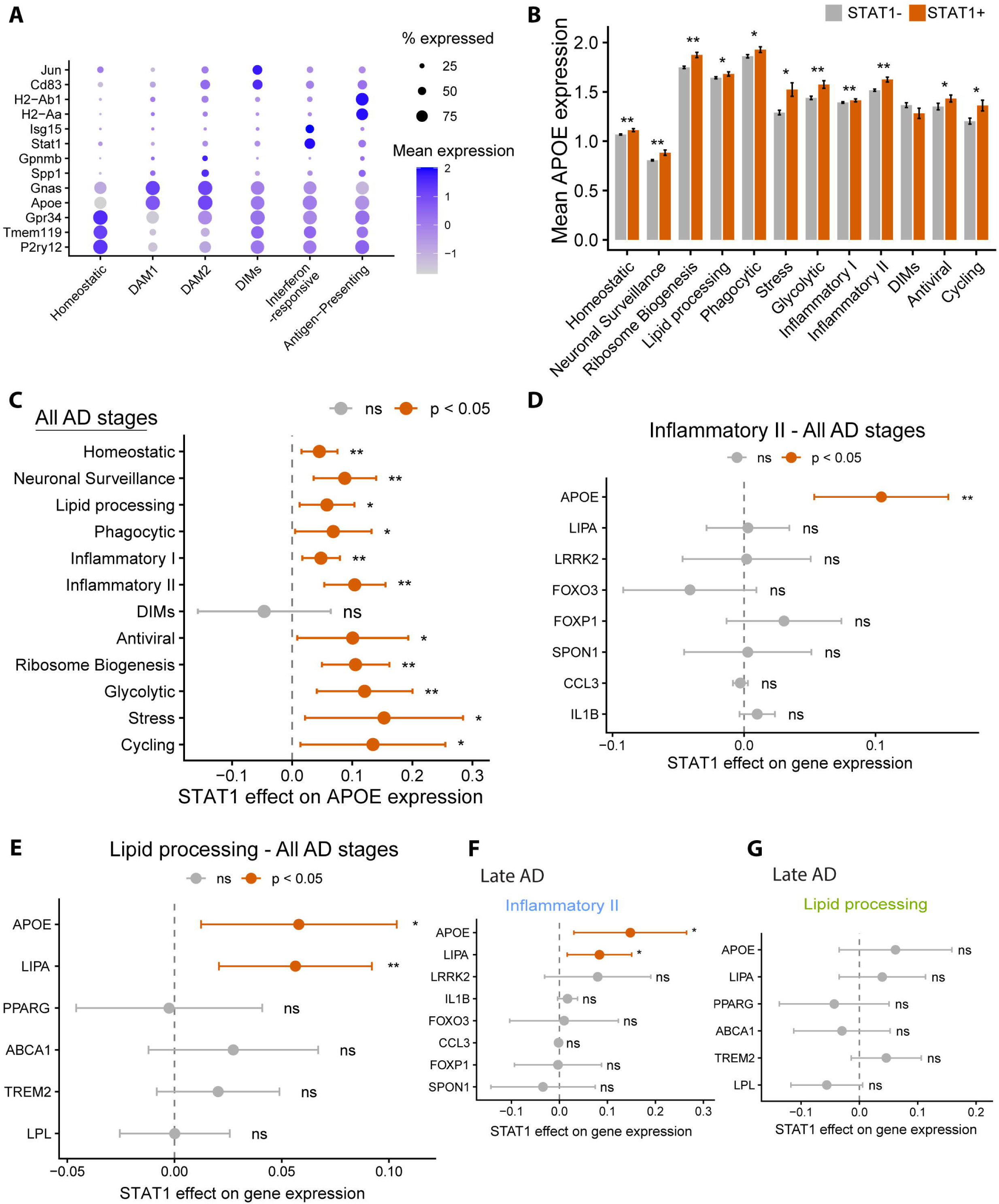
STAT1-associated APOE and lipid-gene expression varies across human AD microglial states and disease stages. **(A)** Dot plot of representative marker genes across homeostatic, DAM1, DAM2, disease-associated intermediate, interferon-responsive, and antigen-presenting microglial states in the mouse AD dataset shown in Fig. 3A. Dot size indicates the percentage of cells expressing each gene, and color indicates scaled mean expression. **(B)** Mean APOE expression in cells with undetected or detected STAT1 transcript across the indicated human AD microglial states. Bars indicate mean expression stratified by STAT1 detection status. *P < 0.05; **P < 0.01. **(C)** State-resolved association between STAT1 detection and APOE expression across all AD stages combined. Points indicate estimated associations with STAT1 detection and horizontal lines indicate 95% confidence intervals. Positive estimates indicate higher APOE expression in cells with detectable STAT1 transcript. Significant associations are highlighted. ns, not significant; *P < 0.05; **P < 0.01. **(D)** Association between STAT1 detection and expression of the indicated genes within Inflammatory II microglia across all AD stages. Points and horizontal lines indicate estimated associations and 95% confidence intervals; positive estimates indicate higher expression in cells with detectable STAT1 transcript. ns, not significant; ***P < 0.001. **(E)** Association between STAT1 detection and expression of the indicated lipid-handling genes within lipid-processing microglia across all AD stages, displayed as in (D). ns, not significant; *P < 0.05; **P < 0.01. **(F)** Association between STAT1 detection and expression of the indicated genes within late-AD Inflammatory II microglia, displayed as in (D). Significant associations are highlighted. ns, not significant; *P < 0.05. **(G)** Association between STAT1 detection and expression of the indicated lipid-handling genes within late-AD lipid-processing microglia, displayed as in (F). ns, not significant.

**Figure S6.**
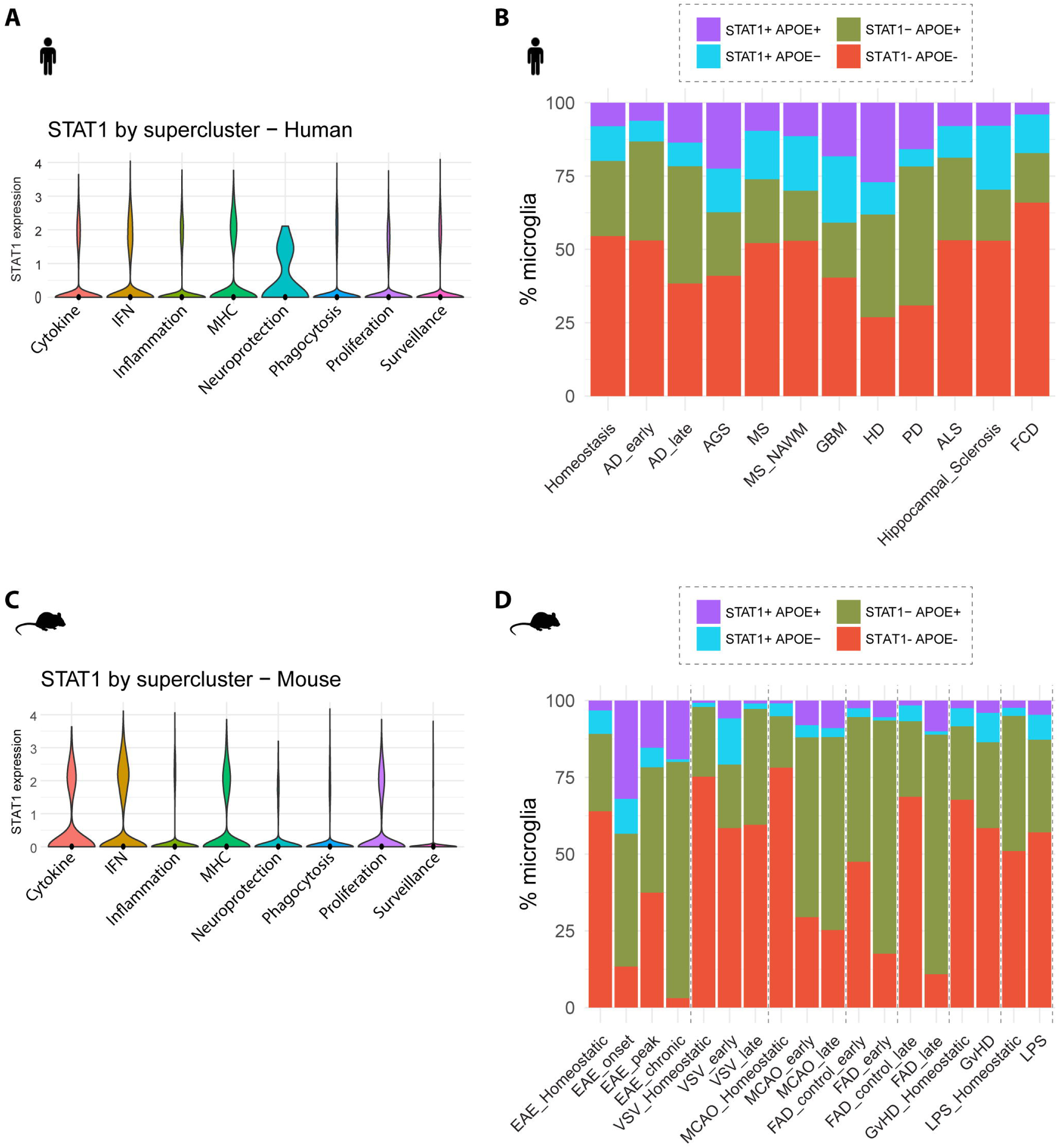
STAT1 expression and STAT1-APOE co-expression recur across diverse human and mouse microglial disease contexts. **(A)** Distribution of STAT1 expression across transcriptionally defined human microglial superclusters, including cytokine-responsive, interferon-associated, inflammatory, MHC-associated, neuroprotective, phagocytic, proliferative, and surveillance programs. Violin plots show the expression distribution within each supercluster. **(B)** Relative proportions of human microglia classified as STAT1 APOE, STAT1 APOE, STAT1 APOE, or STAT1 APOE across aggregated published datasets representing homeostatic and neurological disease contexts, including AD, aging-associated conditions, MS and normal-appearing white matter, glioblastoma, Huntington’s disease, Parkinson’s disease, amyotrophic lateral sclerosis, hippocampal sclerosis, and focal cortical dysplasia. **(C)** Distribution of *Stat1* expression across the corresponding transcriptionally defined mouse microglial superclusters, displayed as in (A). **(D)** Relative proportions of mouse microglia classified according to detectable *Stat1* and *Apoe* transcripts across published datasets encompassing homeostatic and experimentally induced neuroinflammatory, infectious, vascular, neurodegenerative, immune-mediated, and LPS-associated conditions. Dashed vertical lines separate datasets or experimental series where indicated.

**Figure S7.**
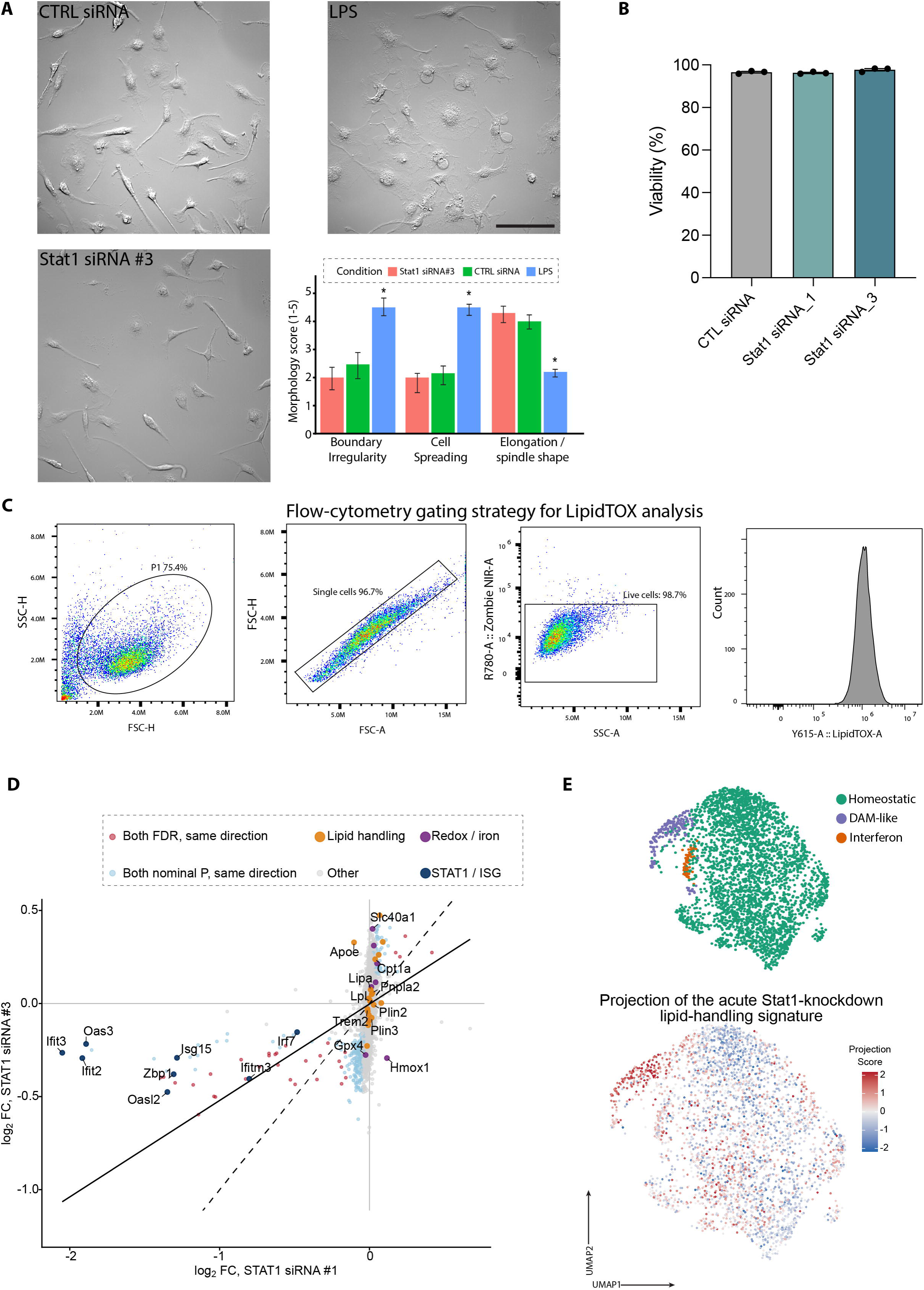
Validation and transcriptional characterization of acute STAT1 depletion in primary microglia. **(A)** Morphological assessment of primary microglia following treatment with control or *Stat1*-targeting siRNA. LPS-treated cultures were included as a positive control for inflammatory activation and morphological responsiveness. Scale bar = 100 µm. *P < 0.05. **(B)** Cell viability following transfection with control or *Stat1*-targeting siRNA. **(C)** Representative flow-cytometry gating strategy used to quantify neutral-lipid LipidTOX fluorescence in primary microglia. Cells were sequentially gated to identify the cellular population, singlets, and viable cells before LipidTOX fluorescence was quantified. The rightmost histogram shows the LipidTOX fluorescence distribution in the analyzed population. **(D)** Comparison of gene-level transcriptional responses to *Stat1* siRNA #1 and #3. Each point represents a gene positioned according to its log fold change relative to control siRNA following siRNA #1 treatment (x-axis) and siRNA #3 treatment (y-axis). Genes showing concordant responses by false-discovery-rate or nominal-P-value criteria are distinguished, together with selected STAT1 and interferon-responsive, lipid-handling, and redox or iron-associated genes; selected genes are labeled. The solid line indicates the fitted relationship and the dashed diagonal indicates equal effects between the two siRNAs. **(E)** Projection of the acute *Stat1*-knockdown lipid-handling transcriptional response onto the adult microglial landscape *in vivo*. Top, UMAP showing homeostatic, DAM-like, and interferon-responsive microglial states. Bottom, per-cell projection scores indicating similarity to the lipid-handling component of the acute knockdown response; positive and negative scores indicate similarity to opposing directions of the projected signature.

**Figure S8.**
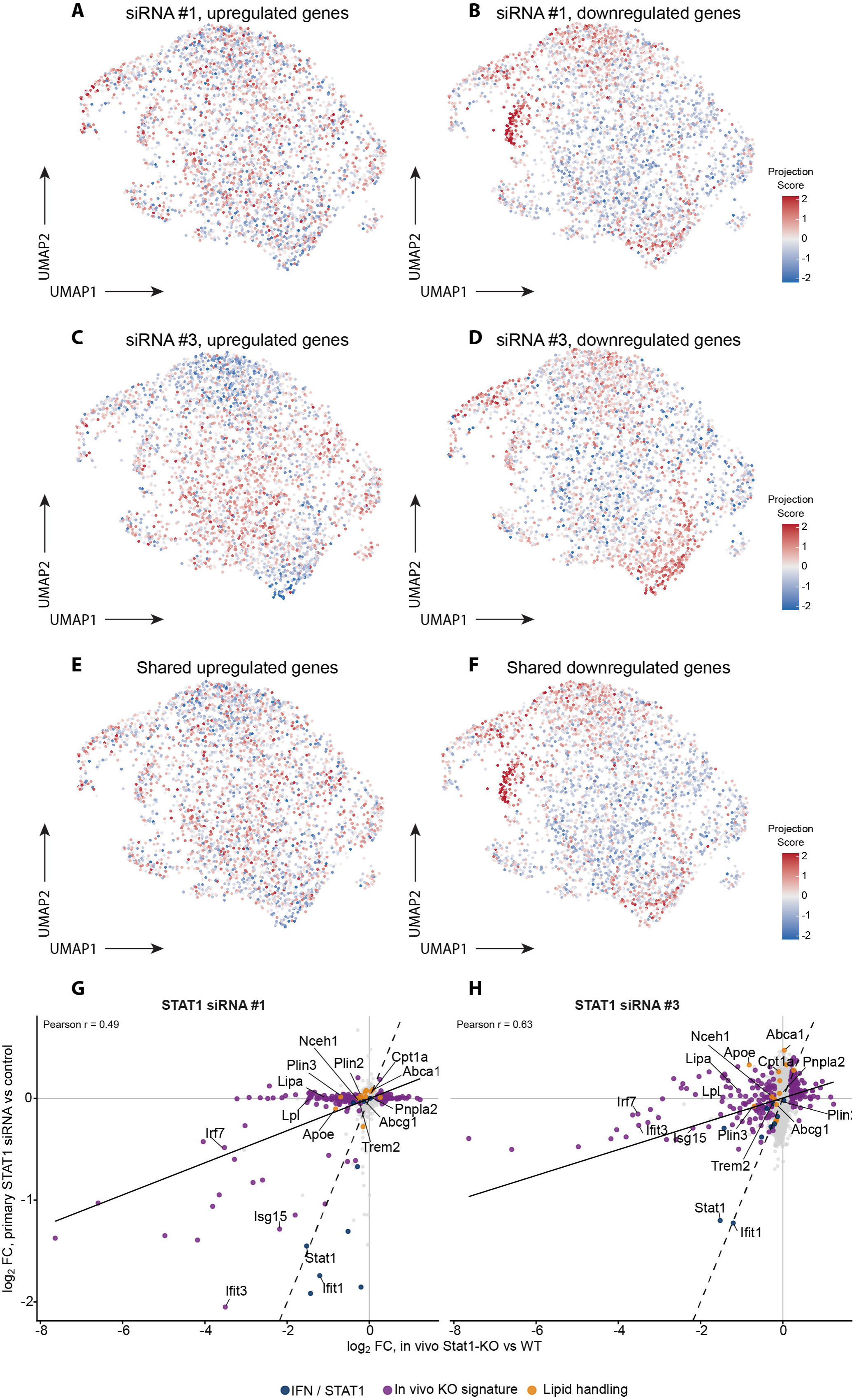
Acute STAT1 depletion reproduces core components of the *in vivo Stat1*-loss response with context-dependent lipid-gene effects. **(A,B)** Projection of genes expressed at higher (A) or lower (B) levels following *Stat1* siRNA #1 treatment onto the adult microglial UMAP *in vivo*. Projection scores indicate the relative representation of each acute-knockdown gene signature across the in vivo microglial landscape. **(C, D)** Projection of genes expressed at higher (C) or lower (D) levels following *Stat1* siRNA #3 treatment onto the same *in vivo* landscape. **(E, F)** Projection of genes showing a shared increase (E) or shared decrease (F) following treatment with *Stat1* siRNA #1 and #3. Color indicates projection score, with positive and negative values representing opposing directions of each transcriptional signature. **(G, H)** Gene-level comparison of transcriptional effects following constitutive *Stat1* loss in adult microglia *in vivo* and acute *Stat1* depletion in primary microglia, using siRNA #1 (G) or siRNA #3 (H). Each point represents a gene positioned according to its log fold change in *Stat1*−/− versus WT microglia *in vivo* (x-axis) and *Stat1* siRNA versus control siRNA in primary microglia (y-axis). Selected STAT1 and interferon-responsive and lipid-handling genes are highlighted and labeled. Gene-level effects are positively correlated between the two models (Pearson r = 0.49 for siRNA #1; r = 0.63 for siRNA #3). Colors distinguish canonical IFN/STAT1 genes, genes belonging to the in vivo *Stat1*-loss signature, and lipid-handling genes. The solid line indicates the fitted relationship and the dashed diagonal indicates equal effect sizes.

**Figure S9.**
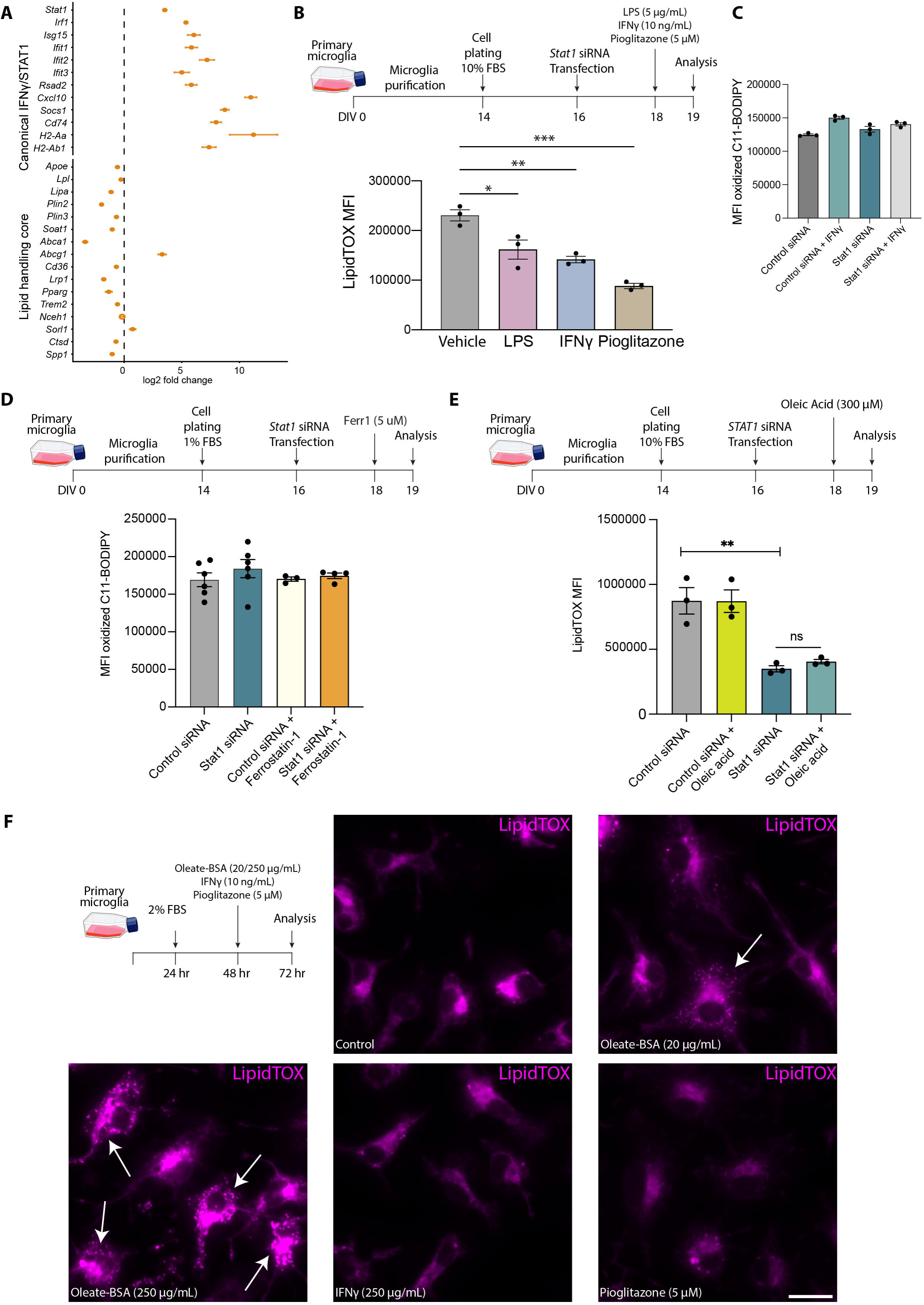
Characterization of lipid peroxidation, neutral-lipid content, and oleate responses following STAT1 depletion in primary microglia. **(A)** Transcriptional response of primary microglia to acute IFNγ stimulation. Forest plot of log_2_ fold changes in selected canonical IFN/STAT1-response genes and core lipid-handling genes following IFNγ treatment of control cells. **(B)** Experimental timeline (top) and LipidTOX median fluorescence intensity (MFI) following inflammatory or metabolic stimulation (bottom). Primary microglia were plated at DIV14, transfected with control or *Stat1*-targeting siRNA at DIV16, treated with vehicle, LPS, IFNγ, or pioglitazone at DIV18, and analyzed by flow cytometry at DIV19. Points represent independent biological replicates; bars indicate mean ± s.e.m. *P < 0.05; **P < 0.01; ***P < 0.001. **(C)** Lipid-peroxidation-associated fluorescence assessed by oxidized C11-BODIPY staining and flow cytometry. Bars indicate C11-BODIPY MFI in control- and *Stat1*-siRNA-treated cells in the presence or absence of IFNγ. Points represent independent biological replicates; bars indicate mean ± s.e.m. **(D)** Experimental timeline (top) and oxidized C11-BODIPY fluorescence measured by flow cytometry (bottom). Primary microglia were purified at DIV0, plated at DIV14, transfected with control or *Stat1*-targeting siRNA at DIV16, treated with or without ferrostatin-1 at DIV18, and analyzed at DIV19. Points represent independent biological replicates; bars indicate mean ± s.e.m. **(E)** Experimental timeline (top) and LipidTOX MFI following oleate supplementation in control and *Stat1*-depleted primary microglia under serum-replete conditions (bottom). Microglia were transfected with control or *Stat1*-targeting siRNA at DIV16, treated with oleate at DIV18, and analyzed at DIV19. Points represent independent biological replicates; bars indicate mean ± s.e.m. **P < 0.01; ns, not significant. **(F)** Representative images showing neutral-lipid loading following oleate supplementation under serum-starved conditions. Arrows indicate lipid-accumulating microglia. Scale bar = 20 µm.

**Figure S10.**
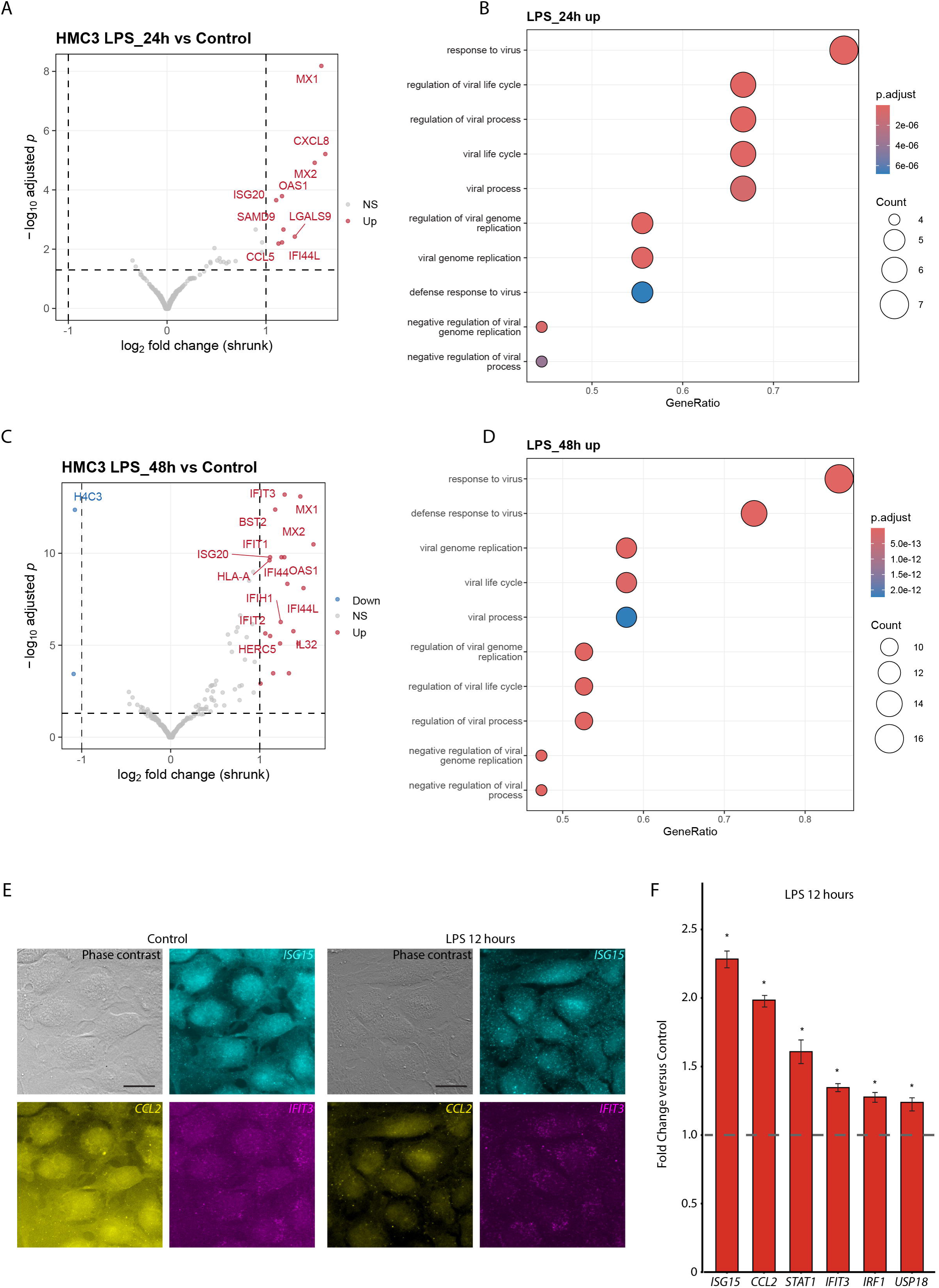
LPS induces interferon- and antiviral-associated transcription in HMC3 cells. **(A)** Volcano plot of differential gene expression in HMC3 cells following 24 h of LPS treatment relative to untreated control cells. The x-axis indicates shrunken log₂ fold change and the y-axis −log_10_ adjusted P value. Differentially expressed genes meeting the applied thresholds are highlighted and selected LPS-responsive genes are labeled. Dashed lines indicate the fold-change and significance thresholds. **(B)** Gene Ontology enrichment analysis of genes expressed at higher levels following 24 h of LPS treatment. The x-axis indicates gene ratio, bubble size the number of genes contributing to each term, and color the adjusted P value. **(C)** Volcano plot of differential gene expression following 48 h of LPS treatment relative to untreated control cells, displayed as in (A). Selected interferon- and antiviral-response genes are labeled. **(D)** Gene Ontology enrichment analysis of genes expressed at higher levels following 48 h of LPS treatment, displayed as in (B). **(E, F)** Representative single-molecule fluorescence *in situ* hybridization images (E) and corresponding quantification (F) validating selected LPS-responsive transcripts in HMC3 cells. Scale bar = 20 µm.

**Figure S11.**
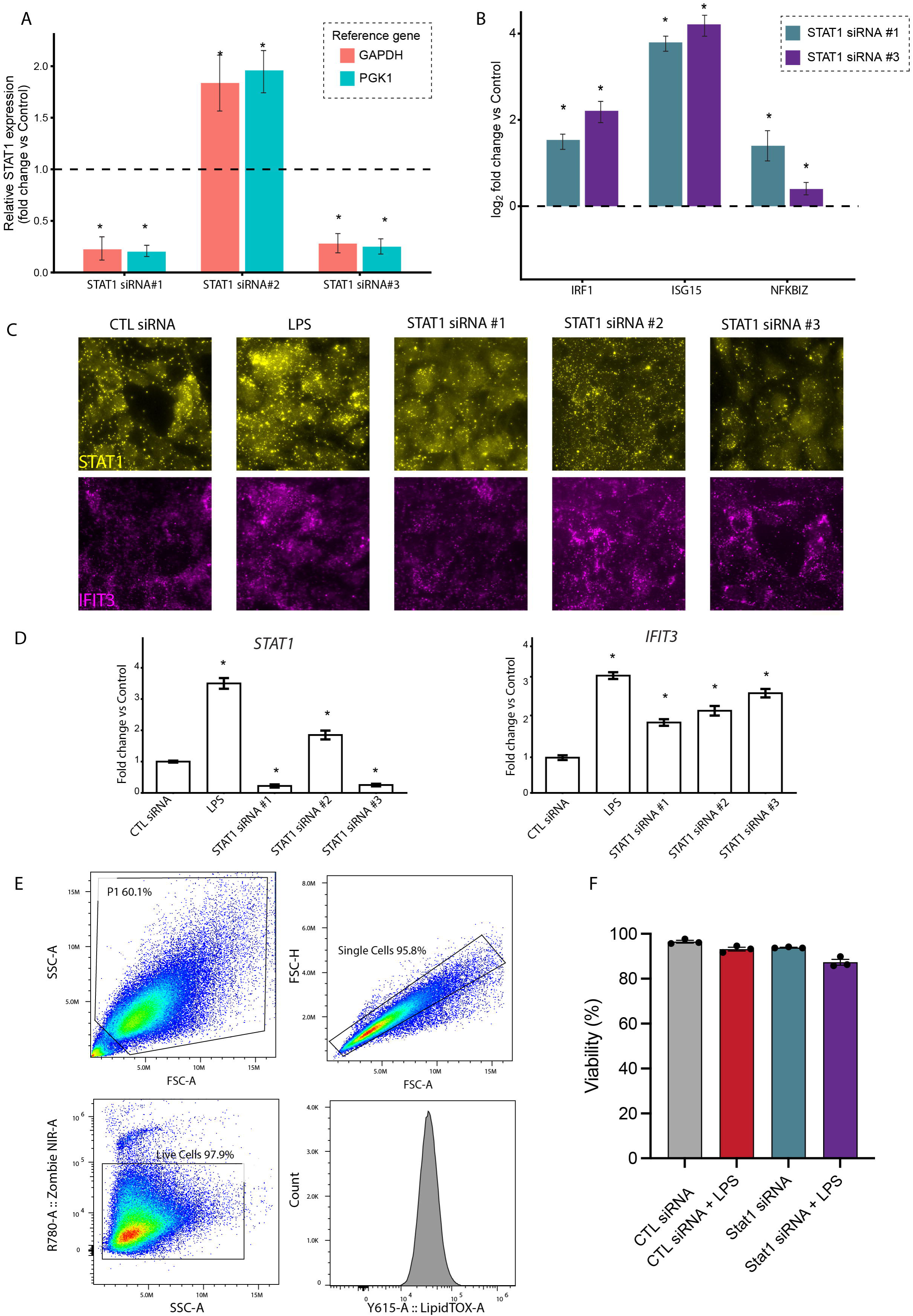
Validation of STAT1 depletion and associated transcriptional and flow-cytometric responses in HMC3 cells. **(A)** Screening of three independent STAT1-targeting siRNAs by quantitative PCR. Relative STAT1 expression is shown as fold change compared with control-transfected cells after normalization to *GAPDH* or *PGK1*. The dashed horizontal line indicates expression in the control condition. Points represent independent replicates; bars indicate mean ± s.e.m. *P < 0.05. **(B)** Expression of selected inflammatory and lipid-handling genes following treatment with STAT1 siRNA #1 or #3. Bars indicate log fold change relative to control-transfected cells for *IRF1, ISG15*, and *NFKBIZ*; positive values indicate higher and negative values lower expression following STAT1 depletion. Error bars indicate the standard error of the shrunken estimate. *P < 0.05. **(C, D)** Representative single-molecule fluorescence *in situ* hybridization images (C) and corresponding quantification (D) of selected transcripts following STAT1 depletion in HMC3 cells. Scale bar = 30 µm. *P < 0.05. **(E)** Representative flow-cytometry gating strategy used to quantify neutral-lipid LipidTOX fluorescence in HMC3 cells. Cells were sequentially gated to identify the cellular population, singlets, and viable cells before LipidTOX fluorescence was quantified. **(F)** Cell viability and LipidTOX-positive-cell measurements across control and STAT1-depleted HMC3 conditions. Points represent independent replicates; bars indicate mean ± s.e.m.

**Figure S12.**
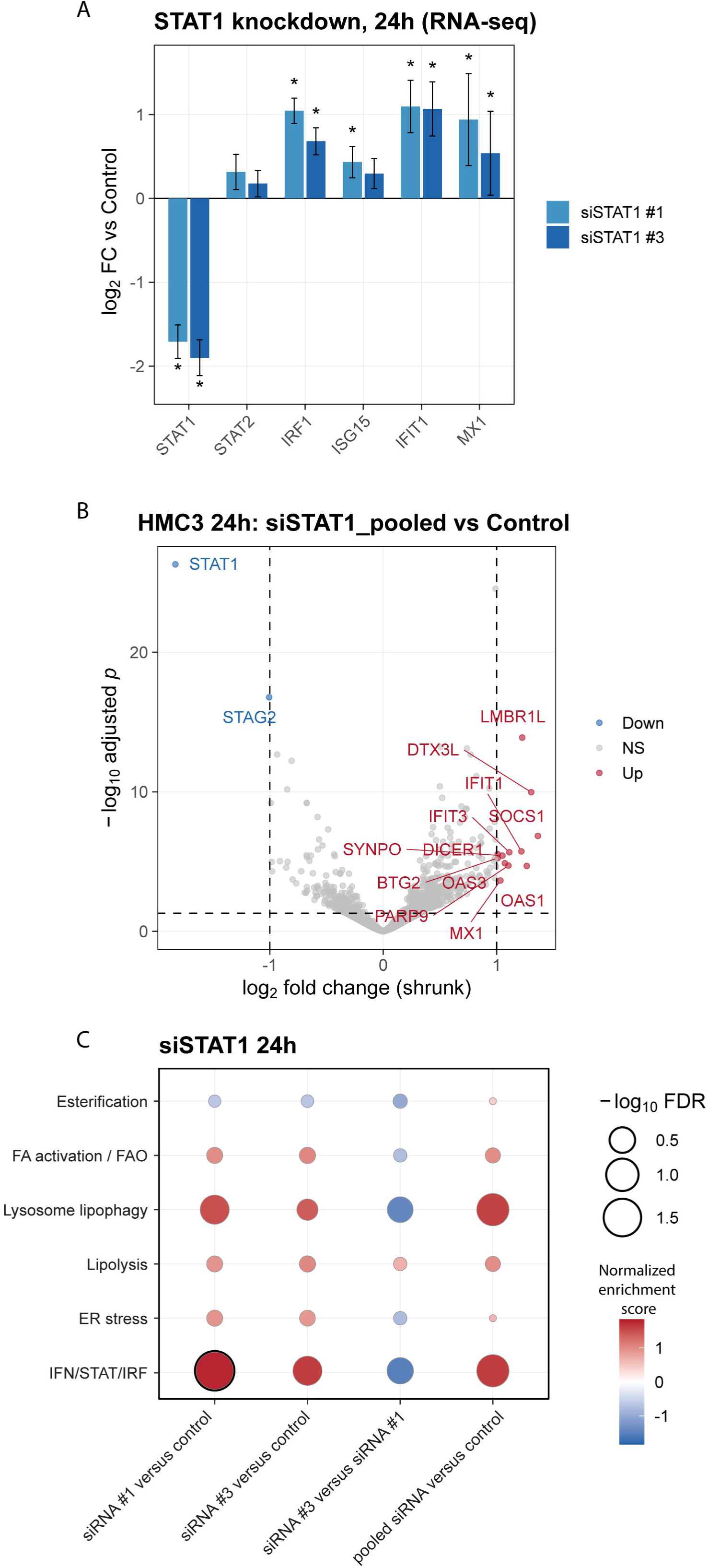
Acute STAT1 depletion in HMC3 cells elicits a secondary inflammatory response and reorganizes lipid-associated programs. **(A)** RNA-seq-derived expression changes in selected STAT1 and interferon-associated genes 24 h after treatment with STAT1 siRNA #1 or #3. Bars indicate log fold change relative to control-transfected cells for *STAT1, STAT2, IRF1, ISG15, IFIT1*, and *MX1*. Error bars indicate the standard error of the shrunken estimate, and asterisks indicate significance in the differential-expression analysis. **(B)** Volcano plot of differential gene expression 24 h after STAT1 depletion in the pooled STAT1-siRNA analysis relative to control. The x-axis indicates shrunken log₂ fold change and the y-axis −log_10_ adjusted P value. Significantly increased and decreased genes are distinguished and selected transcripts labeled. Dashed lines indicate the fold-change and significance thresholds. **(C)** Gene-set enrichment analysis of transcriptional responses 24 h after STAT1 depletion. Columns show STAT1 siRNA #1 versus control, STAT1 siRNA #3 versus control, siRNA #3 versus siRNA #1, and pooled STAT1 siRNA versus control. Rows indicate IFN/STAT1/IRF, endoplasmic-reticulum stress, lipolysis, lysosome and lipophagy, fatty-acid activation and oxidation, and esterification programs. Color indicates normalized enrichment score; bubble size indicates −log_10_ false-discovery rate (FDR).

**Figure S13.**
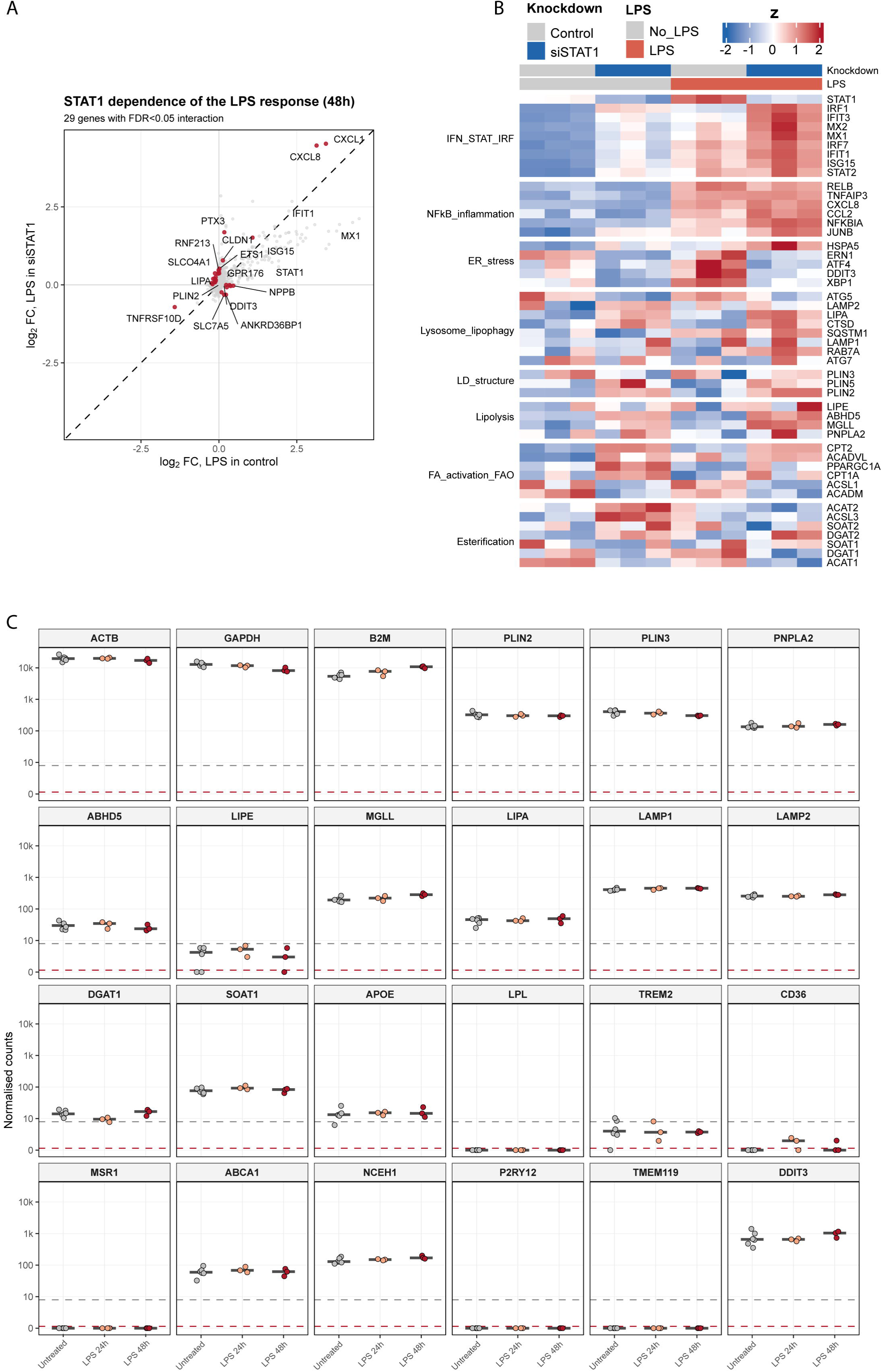
STAT1 depletion modifies selected components of the LPS response, and HMC3 cells incompletely express the microglial lipid-trafficking network. **(A)** Comparison of the 48-h LPS transcriptional response in control and STAT1-depleted HMC3 cells. Each point represents a gene positioned according to its log fold change following LPS treatment in control cells (x-axis) and STAT1-depleted cells (y-axis). The dashed diagonal indicates equal LPS responses in both conditions. Twenty-nine genes show a significant STAT1-by-LPS interaction at false-discovery rate (FDR) < 0.05. Selected interaction-associated inflammatory, stress-associated, and lipid-related genes are labeled. **(B)** Heatmap of expression of representative genes grouped into IFN/STAT1/IRF, NF-κB and inflammatory, endoplasmic-reticulum stress, lysosome and lipophagy, lipid-droplet structure, lipolysis, fatty-acid activation and oxidation, and esterification programs, across control and STAT1-depleted HMC3 cells with or without LPS. Top annotation bars indicate knockdown and LPS-treatment status. Colors indicate row-scaled z-scores. **(C)** Normalized RNA-seq counts for representative housekeeping, lipid-droplet, lipolysis, lysosomal, lipid-trafficking, microglial-identity, and stress-response genes in untreated HMC3 cells and following 24 or 48 h of LPS treatment. Points represent replicate measurements; horizontal bars indicate the mean.

**Figure S14.**
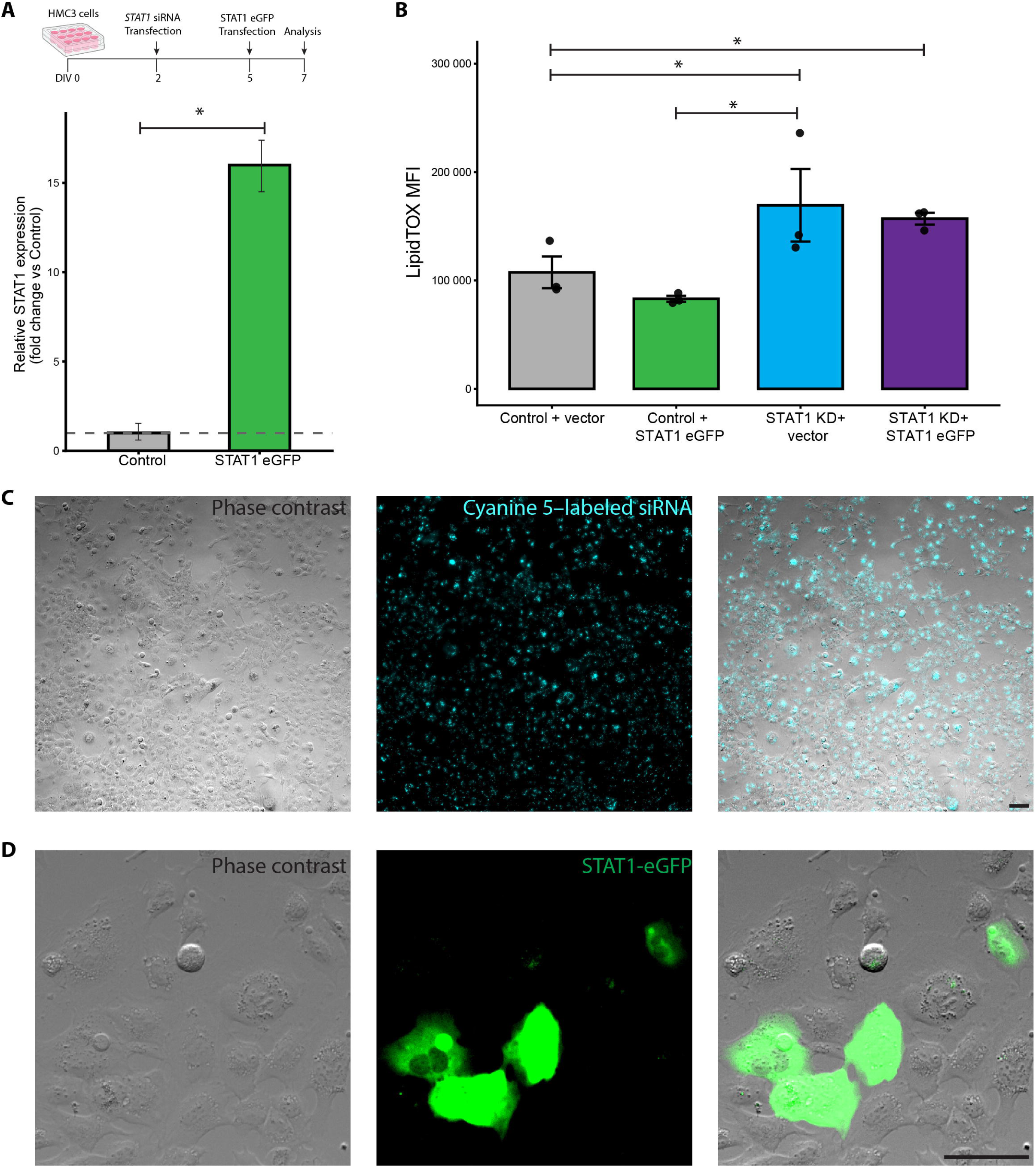
Validation of STAT1 re-expression and analysis of neutral-lipid content in HMC3 cells. **(A)** Experimental timeline (top) and relative STAT1 expression following STAT1-eGFP transfection (bottom). HMC3 cells were plated at DIV0, transfected with STAT1-targeting siRNA at DIV2, transfected with STAT1-eGFP or control vector at DIV5, and analyzed at DIV7. The dashed horizontal line indicates expression in the control condition. Bars indicate mean ± s.e.m. *P < 0.05. **(B)** Neutral-lipid content measured by LipidTOX median fluorescence intensity (MFI) across STAT1-knockdown and STAT1-eGFP re-expression conditions. Points represent independent biological replicates; bars indicate mean ± s.e.m. *P < 0.05. **(C)** Representative phase-contrast, fluorescence (Cyanine 5 channel), and merged overlay micrographs of HMC3 cells transfected with Cyanine 5-labelled siRNA Universal Negative Control. Scale bar = 100 µm. **(D)** Representative phase-contrast, GFP fluorescence, and merged overlay micrographs of HMC3 cells transfected with STAT1-eGFP. Scale bar = 100 µm.

